# *Lmna* deficiency promotes EPHX2 nuclear translocation to ameliorate cardiac dysfunction in mice

**DOI:** 10.64898/2026.02.09.704693

**Authors:** Congting Guo, Ze Wang, Jin Liu, Chenyang Wu, Yuhan Yang, Zhengyuan Lv, Gonglie Chen, Yueshen Sun, Ruilian Bai, Weiyan Sun, Tian Lu, Kai Wang, Zhuang Tian, Xu Zhang, Dongyu Zhao, Shuyang Zhang, Yuxuan Guo

## Abstract

**Background:** Cardiovascular diseases are often associated with altered protein subcellular localization. As a major cause of inherited cardiomyopathy, *LMNA* deficiency could trigger nuclear envelope rupture and broadly impair the localization of nuclear and cytoplasmic proteins. Systemic approaches to identify, dissect and manipulate the localization of endogenous proteins are important for mechanistic and therapeutic investigation.

**Method:** Proximity proteomics of the nuclear lamina was performed specifically in cardiomyocytes in *Lmna*-deficient murine models. AAV-mediated Cas9-based gene silencing and subcellular gene upregulation were conducted via the nuclear localization signal (NLS) and nuclear export signal (NES). Cas9-based somatic mutagenesis was supplemented with the single-strand DNA templates of AAV to achieve robust homology-directed repair (HDR) and targeted NLS knock-in, which translocated cytoplasmic proteins into nuclei.

**Result:** In vivo proximity proteomics detected increased epoxide hydrolase 2 (EPHX2) in cardiomyocyte nuclei in mice carrying germline or cardiac-specific *Lmna* truncating variants. This phenotype was associated with ruptured nuclear envelope. Cas9-mediated *Ephx2* knockout in cardiomyocytes ameliorated cardiac dysfunction in *Lmna*-deficient mice. Strikingly, overexpression of NLS-EPHX2, but not NES-EPHX2, also mitigated cardiac dysfunction. The cardiac protective EPHX2 substrates, epoxyeicosatrienoic acids (EETs), did not alter upon NLS-EPHX2 overexpression. By contrast, the *Lmna-*related DNA damage marker γ-H2AX was reduced. The EPHX-D333A mutant lacking hydrolase activity recapitulated the effects of wildtype EPHX2 in nuclei. AAV-Cas9-based HDR achieved efficient NLS knock-in and EPHX2 nuclear translocation in more than 60% cardiomyocytes, which improved cardiac function.

**Conclusion:** *Lmna* deficiency leads to the nuclear translocation of EPHX2, which ameliorated cardiac dysfunction in a hydrolase-independent manner. AAV-HDR-mediated somatic gene editing provides an efficient approach to manipulate the subcellular localization of endogenous proteins in cardiomyocytes in vivo.

**What is Known?:**

- Cardiovascular diseases are regulated by the changes in protein subcellular localization. In particular, *LMNA*-related cardiomyopathy is associated with nuclear rupture and impaired separation between nuclear and cytoplasmic proteins.
- Epoxide hydrolase 2 (EPHX2) is a cytoplasmic hydrolase that catalyzes the hydrolysis of cardioprotective epoxyeicosatrienoic acids (EETs) and aggravates an array of heart diseases including myocardial infarction and heart failure.
- CRISPR/Cas9-mediated homology-directed repair (HDR) exhibits uniquely high gene editing efficiency in cardiomyocytes with an AAV-based DNA donor, which is suitable for somatic genetic knock-in of small DNA fragments in vivo.

**What New Information Does This Article Contribute?:**

- Lamin-targeted proximity proteomics specifically in cardiomyocytes uncovers novel proteins undergoing subcellular localization changes upon *Lmna* deficiency.
- *Lmna* deficiency leads to EPHX2 nuclear translocation that ameliorates cardiac dysfunction by both mechanisms of cytoplasmic reduction and nuclear induction.
- AAV-HDR-mediated knock-in provides a robust platform to manipulate subcellular localization of endogenous proteins specifically in cardiomyocytes in vivo.

Cardiovascular diseases are often associated with altered protein subcellular localization, but systemic approaches to identify, study and manipulate subcellular localization remain incomplete. This study established an in vivo proximity proteomics approach to identify novel proteins undergoing localization changes relative to the nuclear lamina. In murine models of *LMNA*-related cardiomyopathy, this approach uncovered EPHX2, a classic cytoplasmic hydrolase that aggravates cardiovascular diseases, as a new protein that translocated into cell nucleus and exerted a cardiac protective effect. CRISPR/Cas9-based cardiomyocyte gene editing with an AAV-based DNA donor efficiently achieved NLS knock-in into the *Ephx2* gene, promoted EPHX2 protein nuclear translocation and mitigated cardiac dysfunction with *Lmna* deficiency. These findings indicated a novel avenue to identify and manipulate the subcellular localization changes of endogenous proteins for basic and translational cardiology.

## Introduction

Biomolecules are intricately distributed among subcellular compartments to fulfil their physiological functions. Perturbation of subcellular compartmentation or their localization signals profoundly contribute to pathogenesis[1–4]. Alteration of subcellular localization naturally involves two parallel mechanisms, namely the loss of functions in the original compartments and the potential acquisition of ectopic functions in the new compartments. However, systemic approaches to identify, investigate and manipulate both arms of mechanisms remain to be established for pathophysiological studies and translational medicine.

*LMNA*-related cardiovascular diseases are uniquely associated with subcellular localization changes. *LMNA* encodes A-type lamins including lamin-A and lamin-C, which are intermediate filament proteins that build the nuclear lamina[5, 6]. Lamin-A/C perturbation could impair perinuclear cytoskeletal force and the stiffness of nuclear envelope that resists the force[7–9]. Consequently, nuclear ruptures are frequently observed in *LMNA*-deficient cells [10–12], compromising the boundary between nuclear and cytoplasmic proteins. An array of proteins was reported to change their nucleocytoplasmic distribution in *LMNA* mutant cardiomyocytes[2, 13–16]. Chromatin could also be extruded to cytoplasm to trigger DNA-associated immunity in *LMNA* mutant cells[10, 17]. BioID/BirA*-based proximity proteomics were originally developed to identify lamin-associated proteins in situ, circumventing the technical difficulty in handling the low solubility of lamin polymers[18, 19]. Yet BioID-based characterization of nucleocytoplasmic redistribution upon lamin-A/C perturbation remain to be established in animal models.

Genetic variation in *LMNA* is one of the leading causes of severe cardiomyopathy in human [20–22]. With the slow progress and recent futility of clinical trials on small-molecule drugs[23], gene therapy emerges as a prospective therapeutic approach. However, the unique biological properties of *LMNA* have complicated gene therapy for this disease. For example, adeno-associated virus (AAV)-based gene supplementation was difficult to simultaneously deliver lamin-A and lamin-C while lamin-A supplementation alone was reported pathogenic in wildtype animals[24]. SUN1-targeted gene suppression recently entered the clinical trial stage, but the effectiveness of this strategy is likely restricted to certain genotypes that predominantly perturb nuclear mechanics but not other mechanisms[25, 26]. Correction of *LMNA* mutations by base editing is promising to fully block disease progression[27]. However, with the presence of hundreds of distinct *LMNA* variants[28–32], it is impractical to conduct N-of-1 therapy for each individual variant.

The narrow scope of indication choices and the high cost of canonical gene therapy strategies demand cardiovascular research to explore new therapeutic targets that could cover a broader spectrum of diseases. One potential target is EPHX2 (epoxide hydrolase 2, soluble epoxide hydrolase, or sEH), which is a cytoplasmic enzyme that participates in lipid metabolism by converting epoxides to diols[33]. Inhibition of EPHX2 leads to the accumulation of its protective substrate epoxyeicosatrienoic acids (EETs) [34–36]. Therefore, EPHX2 is considered as a broad-spectrum target for an array of diseases including heart failure[37], myocardial infarction[38, 39], atherosclerosis[40, 41], vascular calcification[42, 43]. Perinuclear localization[44] and hydrolase-independent functions[45] of EPHX2 were also reported, but its impact on *LMNA*-related diseases remains unexplored.

Based on the information above, this study developed an AAV-BioID platform to systemically dissect nuclear lamina-related proteomic changes in models of *LMNA*-related diseases. As the top hit in the BioID analysis, EPHX2 nuclear translocation emerges as a novel target to attenuate *LMNA*-related diseases while holding the potential to alleviate other diseases. An HDR-based gene editing approach is developed to manipulate endogenous EPHX2 nuclear translocation to recapitulate its impact on *LMNA*-related cardiac dysfunction.

## Methods

A list of major resources is available in Supplementary Table 1-5. Additional methods are available in Supplementary Information.

### Animal

Animal strains and procedures were approved by the Institutional Animal Care and Use Committee of Peking University (Animal Protocol #DLASBE0033). *Lmna^Δ/Δ^* mice were generated as previously described[29, 46]. The *Myh6-Cre* mice were purchased from Cyagen Biosciences (strain no. C001041)[47]. The *Lmna^F/F^* mice was acquired from Yixian Zheng lab at Carnegie Institution (The Jackson Laboratory, Strain #026284)[48].

Animals were housed at the Peking University Health Science Center Department of Laboratory Animal Science. All mice were kept in a temperature-controlled room (21 ± 1 °C) with a 12-hour light/dark cycle and had free access to water and normal chow. Littermates of the same sex were randomly assigned to experimental groups. After genotyping at postnatal day 0, neonatal mice under inhalation anesthesia by isoflurane were administrated with recombinant AAVs or vehicle subcutaneously.

### Plasmid

The *lamin-A* and *lamin-B1* coding sequences (CDS) were acquired from murine heart cDNA. The BioID2-HA CDS was from Addgene (#80899)[49]. They were subcloned into the AAV-Tnnt2-Actn2-GFP vector (Addgene #165034)[50] to generate the AAV-Tnnt2-BioID2-HA-Lamin-A and AAV-Tnnt2-BioID2-HA-Lamin-B1 vectors. *Ephx2* sgRNA sequences were subcloned to AAV-U6-sgRNA-Scaffold-*Tnnt2*-SaCas9-HA-OLLAS (Addgene #209781)[51] to generate the AAV-U6-sgRNA-*Tnnt2*-Sacas9-HA vectors. CRISPR/Cas9-AAV9-based somatic mutagenesis (CASAAV) plasmids were produced as previously described[52]. The *Ephx2* CDS was acquired from murine heart cDNA. The D333A point mutation was introduced via primer-directed mutagenesis. They were subcloned into the CMV-3HA-Palmd vector (Addgene # 216833)[53] to generate the CMV-Ephx2 and CMV-Ephx2(D333A) vectors. NLS, NES and mNeonGreen sequences were synthesized by TSINGKE (China). They were subcloned into the AAV-*Tnnt2*-Actn2-GFP vector to generate the AAV-Tnnt2-HA-Ephx2-NLS/NES, AAV-Tnnt2-Ephx2(D333A)-NLS, AAV-Tnnt2-NLS-mNeonGreen, and AAV-Tnnt2-mNeonGreen-Ephx2-NLS vectors. *Ephx2* sgRNA, homology-arm, and *Myl2* promoter sequences were synthesized and subcloned to the AAV-U6-*Palmd* sgRNA1-sgRNA2-*Tnnt2*-Cre-miR122TS vector (Addgene #216830)[53] to generate the AAV-5’HA-NLS-3’HA-*Myl2*-Cre, AAV-sgRNA-U6-Myl2-Cre and AAV-sgRNA-U6-5’HA-NLS-3’HA-Myl2-Cre vectors.

### AAV production

AAV9 was packaged in house or at PackGene Biotech (China) via the three-plasmid method [52, 54]. In brief, AAV9 was packaged in 293T cells with AAV9 Rep-Cap and pHelper (pAd deltaF6, Penn Vector Core). Viruses were then purified on cesium chloride gradients and titered by using the AdEasy adenoviral titer kit (Stratagene). AAV9 titer was determined by quantitative PCR.

### BioID and mass spectrometry

Cardiac BioID in vivo was performed as previously described[53]. The mice were subcutaneously injected with 5×10¹⁰ AAV at P1. For three consecutive days before sample collection, the mice were intraperitoneal injected of 24 mg/kg biotin twice daily. Heart tissues were used for downstream experiments. Biotinylated proteins were enriched using magnetic streptavidin beads (Dynabeads M-280, Invitrogen, 60210) and were washed with a series of SDS buffer, high salt buffer, LiCl buffer and stored in PBS at −80 °C for following LC-MS/MS analysis.

LC-MS/MS analysis was performed at Novogene Corporation. Samples were digested overnight at 37 °C with 200 ng/µL sequencing-grade trypsin. After magnetic bead separation, the supernatant was collected, dried in a speed-vac, and reconstituted in HPLC solvent A (2.5% acetonitrile, 0.1% formic acid). A nano-scale reversed-phase HPLC column was prepared by packing a 100 µm i.d., ∼30 cm fused silica capillary with 2.6 µm C18 beads. Samples were loaded via a Famos autosampler, and peptides were eluted with a solvent B gradient (97.5% acetonitrile, 0.1% formic acid). Eluted peptides were ionized by electrospray and analyzed on an LTQ Orbitrap Velos Elite mass spectrometer. Tandem mass spectra were generated by peptide detection, isolation, and fragmentation. Spectra were searched against a protein database using Sequest (Thermo Fisher Scientific), and data were filtered to a peptide false discovery rate of 1–2%.

### EPHX2 hydrolase activity

HEK293T cells were obtained from Meisen Cell, China (CTCC-DZ-0321). When 70%-80% confluency was achieved, cells were transfected with 2.5 μg EPHX2-expressing plasmids, using 4 μl Lipo8000 (Beyotime, China, C0533) and 125 μl Opti-MEM (Gibco, 31985070) for each well of 6-well plates. After 36h, cells were lysed with 5 μg/ml digitonin at room temperature for 30 min. The supernatant was collected and incubated with 10 μM fluorescent substrate Epoxy Fluor 7. The fluorescence intensity of the product was measured every 3 min at 37 °C and converted into product concentration. Epoxy Fluor 7 can be hydrolyzed by EPHX2 to form 6-methoxy-2-naphthaldehyde, which emits fluorescence at a wavelength of 465 nm upon excitation at 330 nm, thus enabling the monitoring of EPHX2 hydrolase activity.

### EET measurement

EETs were measured by ELISA or UPLC-MS/MS. For ELISA, an EETs ELISA kit (Win-Win Biotech, Shanghai, China, SY-ELA0307) was used. Heart tissues were lysed in normal saline, centrifuged, and the supernatant was diluted prior to assay according to the manufacturer’s protocol. EETs concentrations were calculated from absorbance measured at 450 nm.

The UPLC-MS/MS method was performed as previously described[38]. All performance was conducted on ice. Briefly, ∼50 mg of tissue was homogenized in 500 μL ice-cold methanol containing 5 mM butylated hydroxytoluene (BHT), 10 μL acetic acid, and 1 μL eicosanoid internal standard mixture (6-keto-PGF1α-d4, PGE2-d4, LTB4-d4, 11,12-DHET-d11, 20-HETE-d6, 5-HETE-d8). After centrifugation, 700 μL triple-distilled water and 1 mL ethyl acetate were added to the supernatant. Following vortex and centrifugation, the upper organic phase was evaporated to dryness under nitrogen and reconstituted in 100 μL 30% acetonitrile with 10 min vortexing. and was subjected to Acquity UPLC coupled with a 5500 QTRAP hybrid triple quadrupole-linear ion trap mass spectrometer (AB Sciex) equipped with a turbo electrospray ionization source. Analytes were monitored in negative ion mode using multiple reaction monitoring (MRM). Raw data were processed with Analyst 1.6, and quantification was performed using MultiQuant 3.0.2.

### Statistics

Statistics were performed using Prism (GraphPad 8.0.2) software. Images were analyzed using ImageJ software. Data are presented as mean ± SD. Statistical analysis of all data was determined using the Mann-Whitney *U* test for two-group comparisons and Kruskal-Wallis *H* test for comparison of multiple groups. Kaplan-Meier curves compared with log-rank test were used to evaluate survival. Statistical significance was defined by \**P* < 0.05, \*\**P* < 0.01, \*\*\**P* < 0.001.

### Data availability

Data supporting the findings in this study are available from the corresponding authors on reasonable request. Plasmids are available at Addgene. Sequencing data has been deposited at GSA (https://ngdc.cncb.ac.cn/gsa, CRA036605). Mass spectrometry data are available from supplementary source data.

## Results

### In vivo proximity proteomics identifies ectopic nuclear translocation of EPHX2 in *Lmna*-deficient cardiomyocytes

To establish cardiomyocyte-specific nuclear-lamina proximity labeling approaches, AAV9 was constructed to express BioID2-lamin-A (BioID-LA) and BioID2-lamin-B1 (BioID-LB1) via a cardiomyocyte-specific *Tnnt2* promoter (**Figure 1A**). 5 × 10^10^ AAV9 was subcutaneously injected to postnatal day 1 (P1) mice and 24 mg/kg biotin was intraperitoneally injected twice a day for three days before ventricles were collected at P14 (**Figure 1A**). Cardiac immunofluorescence demonstrated colocalization between streptavidin (SA)-labelled proteins and the HA tag of BioID2-LA/LB1 (**Figure 1B**), or pericentriolar material 1 (PCM1), a marker of cardiomyocyte nuclei (**Figure 1C**). Western blots showed much lower levels of BioID2-LA/B1 than endogenous lamin-A/B1 in the heart (**Figure S1A**). SA pulldown analysis of the biotinylated proteins showed that BioID-LA/LB1 triggered the biotinylation of many proteins with similar molecular weights (**Figure 1D**).

**Figure 1.**
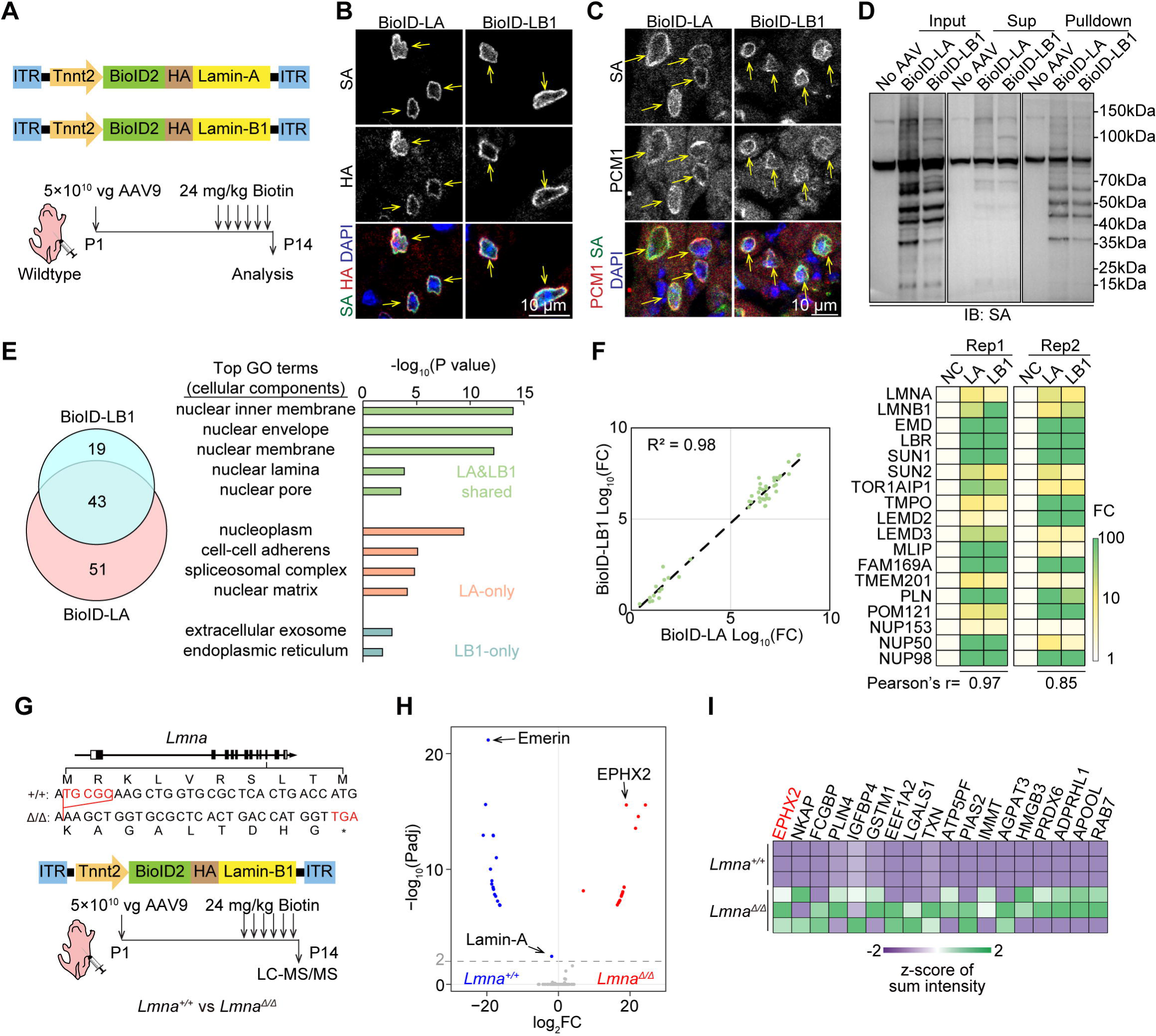
In vivo proximity proteomics reveals EPHX2 enrichment at the nuclear-lamina of *Lmna^Δ/Δ^*cardiomyocytes. **A**, Diagram of AAV vectors and the workflow of nuclear-lamina proximity proteomics. AAV was injected subcutaneously. Biotin was injected intraperitoneally every 12 hours for 3 days before tissue collection. **B** and **C**, Immunofluorescence images of P14 myocardium. Yellow arrows indicate BioID2-labeled nuclei co-stained with HA (**B**) or PCM1 (**C**). **D**, Western blot images of P14 myocardium. **E**, Venn diagram (**left**) and top gene ontology (GO) items (**right**) of proteins identified by BioID-LB1 and BioID-LA. *P* values were generated by Fisher’s exact test corrected by the Bonferroni’s method. **F**, Correlation analysis of BioID-enriched proteins **(left)** and Heatmap of BioID-enriched nuclear lamina-associated proteins **(right)**. R^2^, correlation coefficient. **G**, Schematic of *Lmna^Δ/Δ^* mouse genotype and experimental workflow. **H**, Volcano plot of differential proteomic analysis. *P* values were generated by student’s *t* test corrected by the Bonferroni’s method. **I**, Heatmap of significantly enriched proteins in the *Lmna^Δ/Δ^* heart. ITR, inverted terminal repeat. HA, hemagglutinin. LA, lamin-A. LB1, lamin-B1. SA, streptavidin. Sup, supernatant. IB, immunoblotting. FC, fold change. Rep, replicate. NC, negative control. *Padj*, adjusted *P* value. LC-MS/MS, liquid chromatography-tandem mass spectrometry.

Liquid chromatography-tandem mass spectrometry (LC-MS/MS) was next performed to identify these biotinylated proteins. A threshold of >2 fold changes (FCs) in protein abundance relative to the negative control uncovered a total of 62 and 94 significantly enriched proteins in the BioID-LA and BioID-LB1 groups with 43 shared proteins (**Figure 1E**). Gene ontology (GO) analysis confirmed that nuclear lamina-related terms were significantly enriched among the shared proteins, but not in the BioID-LA or BioID-LB1-specific groups (**Figure 1E**). Moreover, a high correlation between the FCs of the shared proteins was detected between the BioID-LA and BioID-LB1 datasets. Eighteen known nuclear-lamina-related proteins were identified in this shared subset (**Figure 1F**). By contrast, the FCs of proteins identified only in one group exhibited low correlation with those in the other (**Figure S1B**).

Next, AAV-BioID-LB1 was applied to control or *Lmna^Δ/Δ^*mice that carry a homozygous truncating mutation in *Lmna*[14, 46] to investigate the differences in cardiomyocyte-specific nuclear-lamina proteomes in vivo (**Figure 1G**). Western blot confirmed the successfully induction of biotinylation and the mutation of lamin-A/C (**Figure S1C**). LC-MS/MS analysis revealed significantly decreased lamin-A and emerin[14, 55] in the *Lmna^Δ/Δ^* group (**Figure 1H and S1D**) as surrogate markers to validate the successful detection of proteomic alterations in the nuclear lamina of *Lmna^Δ/Δ^* cells. Notably, EPHX2 was the top hit among proteins that were upregulated in the *Lmna^Δ/Δ^* group (**Figure 1H and 1I**). Immunofluorescence of isolated adult cardiomyocytes and embryonic fibroblasts verified low nuclear EPHX2 in control but increased signals in *Lmna^Δ/Δ^* mice (**Figure 2A and 2B**). Nucleocytoplasmic fractionation analysis also indicated reduced cytoplasmic EPHX2 and increased nuclear EPHX2 in *Lmna^Δ/Δ^* hearts (**Figure 2C**). No changes in *Ephx2* mRNA and protein expression were detected by Real-Time quantitative-PCR (RT-qPCR) and western blots (**Figure 2D**).

**Figure 2.**
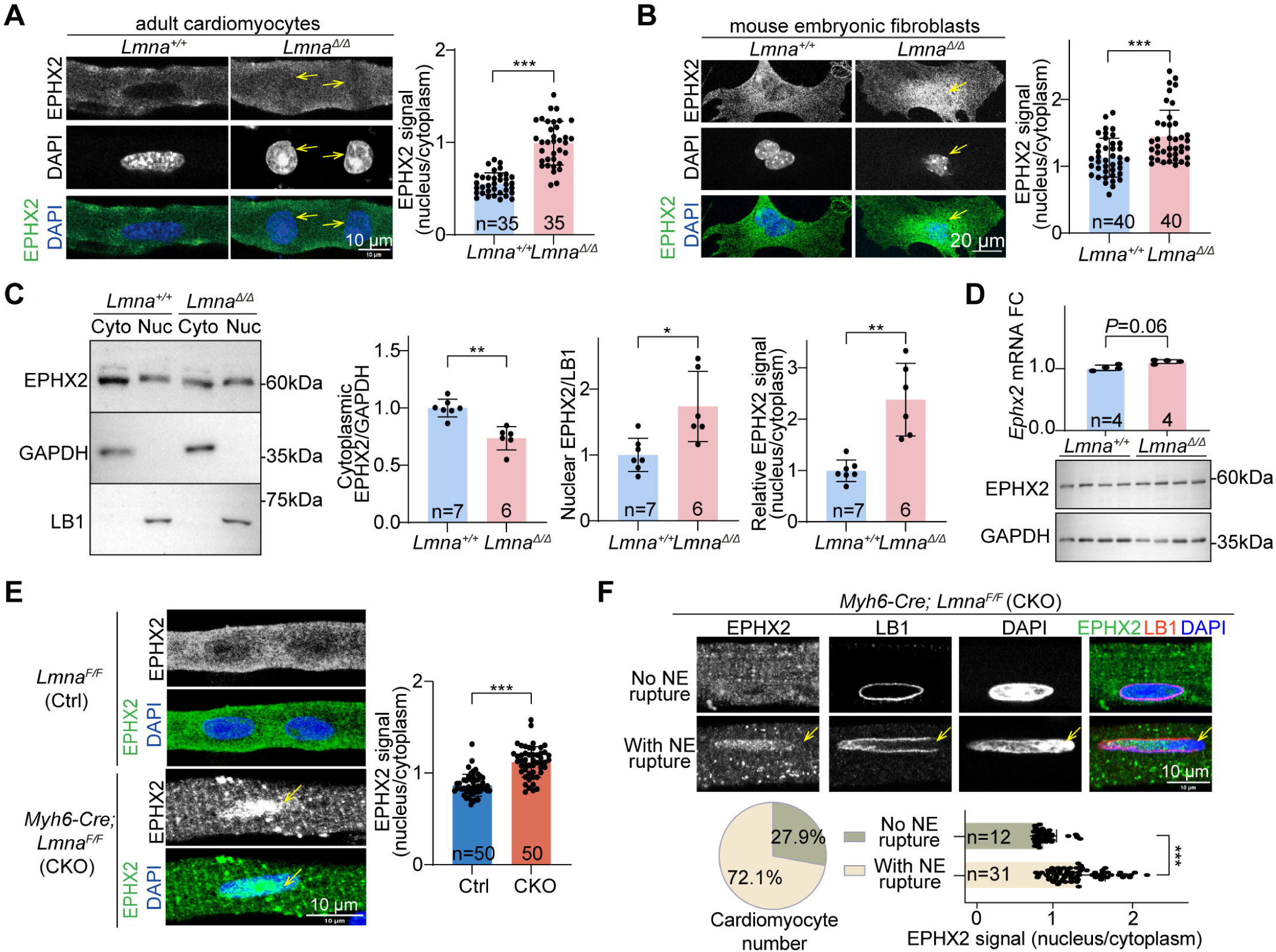
EPHX2 exhibits enhanced nuclear localization in *Lmna^Δ/Δ^* mice. **A** and **B**, Immunofluorescence imaging and quantification of P14 isolated cardiomyocytes (**A**) and E13.5 mouse embryonic fibroblasts (**B**). **C**, Western blot images and quantification of nuclear/cytoplasmic EPHX2 in *Lmna^Δ/Δ^* mouse hearts. **D**, Quantification by qRT-PCR and western blot images of *Lmna^Δ/Δ^* hearts. **E**, Immunofluorescence images and quantification of EPHX2 in isolated cardiomyocytes of P14 CKO mice. **F**, Immunofluorescence images and quantification of NE rupture and EPHX2 location. *P* values were generated by Mann-Whitney *U* test. \**P*<0.05, \*\**P*<0.01, \*\*\**P*<0.001, mean ± SD. FC, fold change. n represents numbers of cells (**A**, **B**, **E** and **F**) or mice (**C** and **D**). LB1, lamin-B1. HA, hemagglutinin. LSL, loxP-stop-loxP. NE, nuclear envelop.

The *Myh6-Cre; Lmna^F/F^* (CKO) mice were next analyzed to confirm EPHX2 subcellular localization changes specifically in cardiomyocytes by immunofluorescence (**Figure 2E**). Notably, DeepLoc 2.1 [56], a protein language model-based subcellular localization predictor, indicated low intrinsic capacity of EPHX2 to enter nuclei (**Figure S1E**). Sequence analysis by WolF PSORT[57], Cell-PLoc[58], cNLS Mapper[59] and NucPred[60] also demonstrated the low likelihood of a nuclear localization signal (NLS). By contrast, a positive correlation between EPHX2 nuclear translocation and incomplete lamin-B1 staining at nuclear periphery was detected (**Figure 2F**), indicating NE rupture-mediated diffusion of EPHX2 from cytoplasm into nuclei.

### CRISPR/Cas9-mediated *Ephx2* ablation alleviates *Lmna*-associated cardiac dysfunction

The impacts of reduced cytoplasmic EPHX2 and increased nuclear EPHX2 on *Lmna*-deficient hearts were separately investigated in the following sections. Firstly, an AAV vector was constructed to achieve SaCas9[51]-mediated *Ephx2* knockout in CKO mice (**Figure 3A**). Two sgRNAs targeting *Ephx2* were validated by next-generation sequencing to trigger >10% DNA insertions and deletions (indels) in the heart (**Figure 3B**). SgRNA1 triggered more efficient depletion of EPHX2 proteins than sgRNA2 (**Figure 3C**) and then was selected for following experiments. The CKO mice exhibited a rapid decline of fractional shortening (FS), an increase of left ventricular inner diameters (LVIDs) and death by three weeks after birth as reported[61] (**Figure 3D-E**). AAV-SaCas9-based *Ephx2* knockout significantly restored these parameters (**Figure 3D-E**). Enzyme-linked immunosorbent assay (ELISA) and mass-spectrometry analysis confirmed elevated EET levels upon *Ephx2* knockout in the heart tissues (**Figure 3F**).

**Figure 3.**
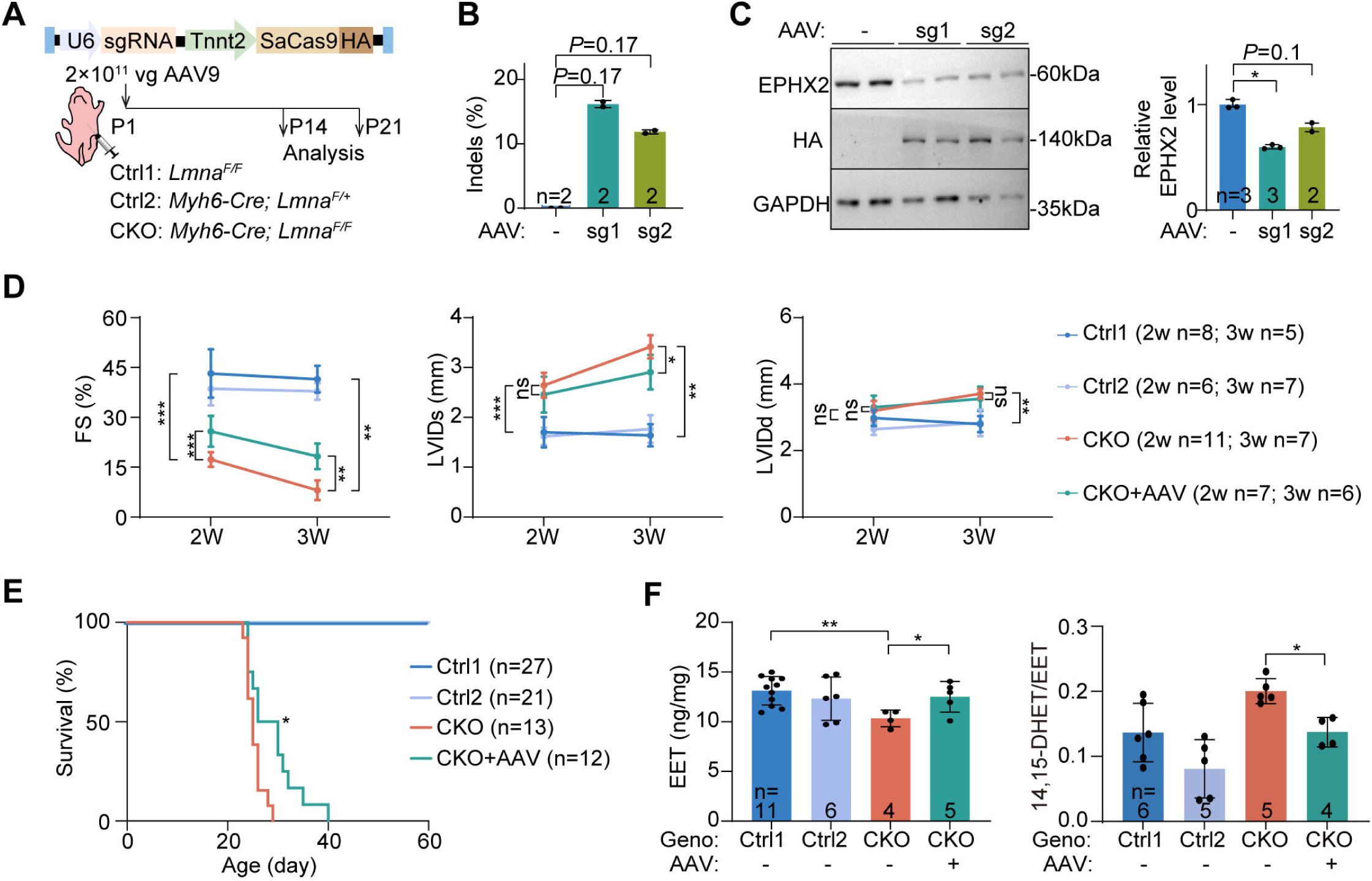
Knocking out *Ephx2* in *Lmna* CKO mice alleviates cardiac dysfunction. **A**, Diagram of AAV vectors and the workflow of knocking out *Ephx2*. AAV was injected subcutaneously. **B**, Editing efficiency of *Ephx2* by amplicon sequencing analysis in hearts. **C**, Western blot analysis and quantification of AAV-treated cardiac tissues. **D**, Echocardiography of cardiac function and dimensions at 2 or 3 weeks after AAV treatment. **E**, Survival curve with the log-rank test between the CKO and CKO+AAV groups, \**P*<0.05. **F**, EET quantification of cardiac tissues by enzyme linked immunosorbent assay (ELISA) and mass-spectrometry analysis. *P* values were generated by Mann-Whitney *U* test. \**P*<0.05, \*\**P*<0.01, \*\*\**P*<0.001, mean ± SD. n represents numbers of mice. FS, fraction shortening. LVIDd, left ventricular internal dimension in diastole. LVIDs, left ventricular internal dimension in systole. EET, epoxyeicosatrienoic acid.

As an orthogonal approach to validate these results, AAV9-*Tnnt2*-Cre (AAV-Cre) or AAV9-*Tnnt2*-Cre-U6-*Ephx2*-sgRNA (AAV-sgRNA) vectors were administered to *Lmna^F/F^*; Rosa^CAG-LSL-SpCas9-GFP^ mice (**Figure S2A**). This CRISPR/Cas9-AAV9-based somatic mutagenesis (CASAAV)[52, 62] strictly limited Cre-based *Lmna* inactivation and sgRNA-based *Ephx2* inactivation in the same cardiomyocytes in mice. Two sets of sgRNAs were validated by western blots and sgRNAs with higher EPHX2 depletion efficiency were selected (**Figure S2B**). AAV transduction rates were confirmed by counting GFP-positive cells in the heart, confirming similar gene delivery efficiency between the groups (**Figure S2C**). Cardiac systolic dysfunction and animal death upon *Lmna* deficiency was rescued by *Ephx2* knockout (**Figure S2D-E**). Enhanced cardiac EET levels were also confirmed (**Figure S2F**).

### The nuclear overexpression of EPHX2 ameliorates *Lmna*-associated cardiac dysfunction

Next, two AAV vectors were constructed to express EPHX2 tagged by a nuclear localization signal (NLS) or a nuclear export signal (NES) to investigate its effect in specific subcellular compartments (**Figure 4A**). Western blots showed successful expression of EPHX2-NLS/NES, but the amount is much lower than the endogenous proteins (**Figure 4B**). We reasoned that this overexpression is adequate to study nuclear EPHX2 because the volume of nuclei is less than 10% of the cytoplasm in cardiomyocytes. Immunofluorescence of heart cryosections confirmed the expected localization of these exogenous EPHX2 (**Figure 4C**). Interestingly, cardiac function was restored in CKO mice that were treated with EPHX2-NLS but not EPHX2-NES, as supported by the increased FS and reduced LVID (**Figure 4D**).

**Figure 4.**
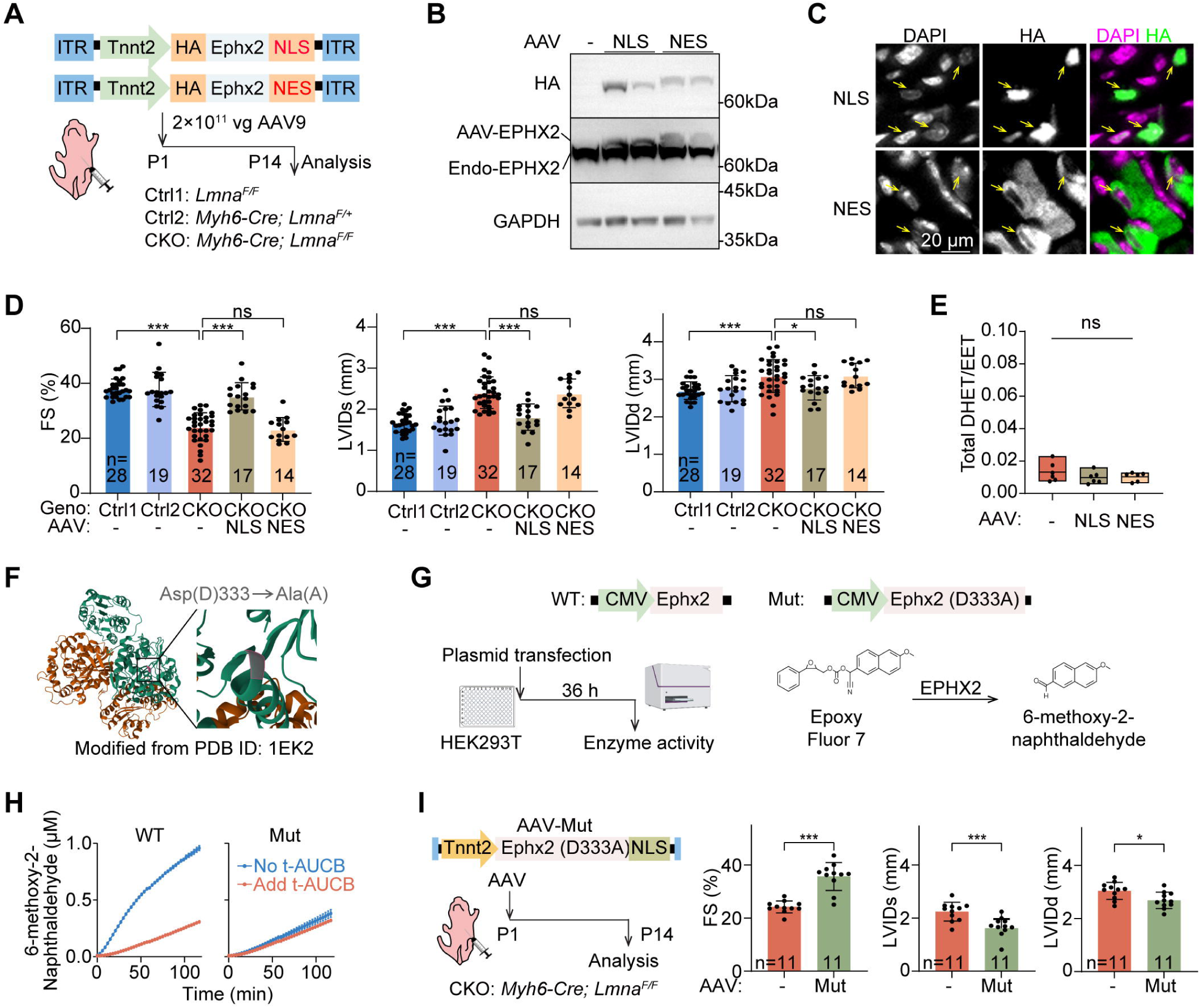
Overexpression of nuclear EPHX2 in *Lmna* CKO mice alleviates cardiac dysfunction. **A**, Diagram of AAV vectors and the workflow of overexpressing nuclear EPHX2. AAV was injected subcutaneously at P1. **B**, Western blot analysis of EPHX2 overexpression. **C**, Immunofluorescence images show the expression and localization of EPHX2 in the myocardium of P14 mouse hearts. The yellow arrow points to the nucleus. **D**, Echocardiography of cardiac function and structure. **E**, Total DHET/EET in hearts by ultra performance liquid chromatography-tandem mass spectrometry (UPLC–MS/MS)-based lipidomic. 6 animals per group. The Kruskal-Wallis H test was used. ns, not significant. **F**, Schematic diagram of inactivating mutation sites in EPHX2 hydrolase. **G**, Work Flow for the detection of EPHX2 hydrolase activity. **H**, Dynamic curve of EPHX2 hydrolysis product concentration. **I**, Schematic diagram of AAV vectors and experimental workflow (left). Echocardiographic analysis of cardiac function in CKO mice (right). CKO mice were injected with 2×10^11^ AAV9 to specifically overexpress hydrolase-deficient EPHX2 in cardiomyocyte nuclei at P1. Echocardiographic analysis was performed at P14. *P* values were generated by Mann-Whitney *U* test. \**P*<0.05, \*\*\**P*<0.001, mean ± SD. n represents numbers of mice. HA, hemagglutinin. NLS, nuclear localization sequence. NES, nuclear export sequence. t-AUCB, trans-4-(4-[3-Adamantan-1-yl-ureido]-cyclohexyloxy)-benzoic acid, a small-molecule inhibitor of EPHX2 hydrolase activity. FS, fraction shortening. LVIDd, left ventricular internal dimension in diastole. LVIDs, left ventricular internal dimension in systole.

EETs were not affected by EPHX2-NLS in the heart, suggesting a hydrolase-independent function of EPHX2 in cell nuclei (**Figure 4E and S3**). To further test this idea, a hydrolase-deficient EPHX2 mutant (EPHX2 D333A) was generated by disrupting the key residue for its enzyme activity (**Figure 4F**)[63]. The lack of hydrolysis activity of EPHX2-D333A was validated in human embryonic kidney 293T (HEK293T) cells via a fluorescence hydrolysis assay (**Figure 4G**) in comparison to wildtype EPHX2 and its specific hydrolase inhibitor t-AUCB (**Figure 4H**). Subsequently, AAV-EPHX2-D333A was administered to CKO mice and echocardiographic analysis confirmed the hydrolase-independent effect of EPHX2 in alleviating *Lmna*-associated cardiac dysfunction (**Figure 4I**).

To further characterize the working mechanism of nuclear EPHX2 specifically in cardiomyocytes, AAV-delivered EPHX2-NLS was tagged with mNeonGreen for fluorescence-activated cell sorting of isolated cardiomyocytes (FACS, **Figure 5A**). Switching mechanism at 5’ end of the RNA transcript-sequencing (SMART-seq) was conducted on the sorted cells for differential expression analysis. Principal component analysis (PCA) revealed a clear separation of control and CKO groups while EPHX2-NLS treatment reduced this difference (**Figure 5B**). 619 downregulated genes and 668 upregulated genes in CKO cardiomyocytes were restored by EPHX2-NLS treatment (**Figure 5C**). Gene ontology analysis demonstrated energy metabolism, cardiac contraction, cell adhesion, inflammatory response and DNA-damage related cell death as the main rescued biological process (**Figure 5C**). Immunofluorescence and western blotting of γH2AX in the heart further confirmed a new role of EPHX2-NLS in reducing DNA damages (**Figure 5D-E**).

**Figure 5.**
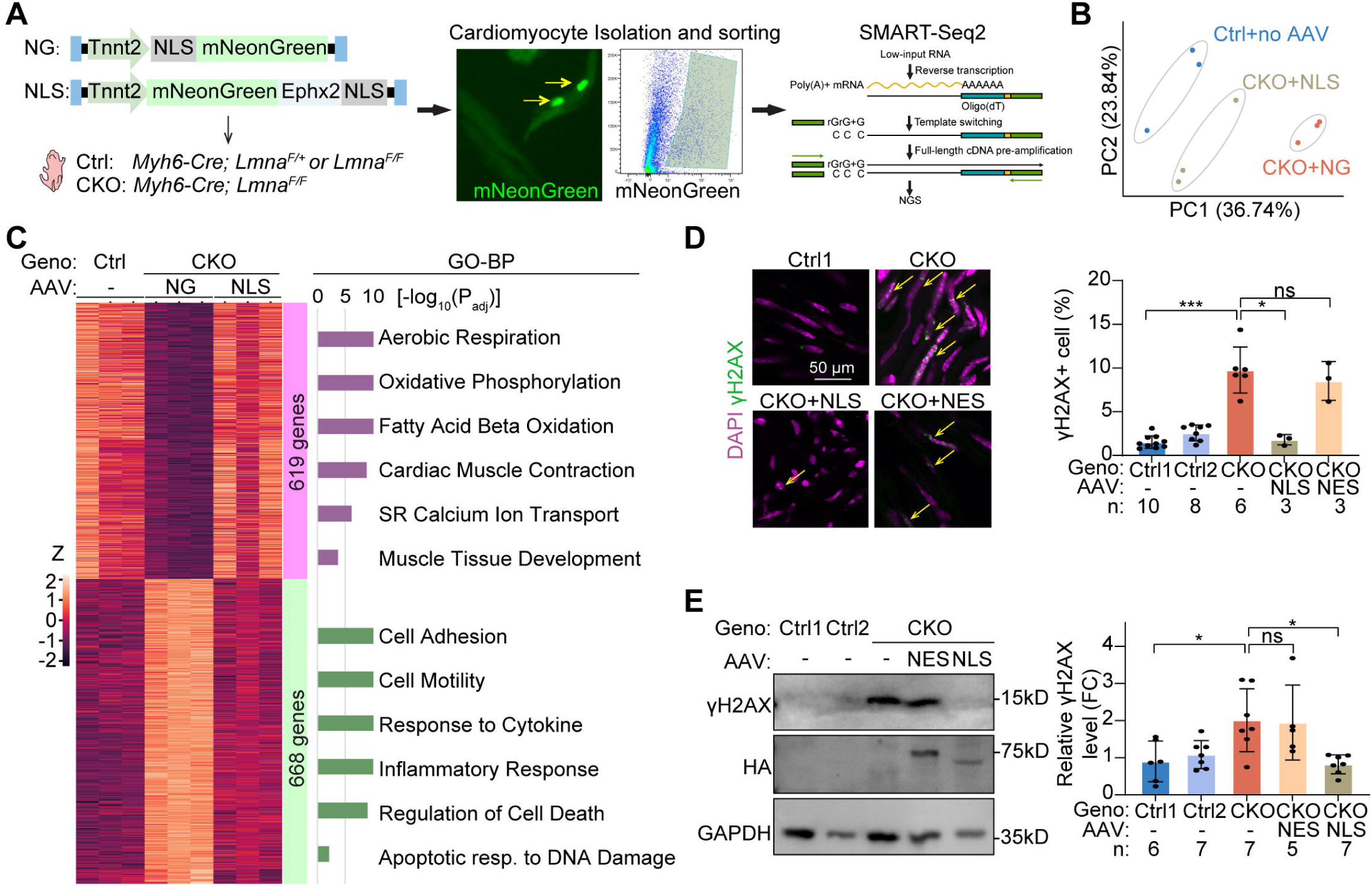
Nuclear EPHX2 overexpression restores cardiac function in a hydrolase-independent manner. **A**, Diagram of AAV vectors and the workflow of SMART-seq2. **B**, PCA plot of the transcriptome sequencing results. **C**, Heatmap of differentially expressed genes and Gene Ontology Biological Process (GOBP) analysis of genes rescued by AAV-NLS. *P*-value indicates a modified Fisher’s Exact *P* value. **D**, Immunofluorescence imaging of γH2AX in P14 mouse hearts. **E**, Western blot imaging and quantification of γH2AX in P14 hearts. *P* values were generated by Mann-Whitney *U* test. \**P*<0.05, \*\*\**P*<0.001, mean ± SD. n represents numbers of mice. SMART-seq, switching mechanism at 5’ end of the RNA transcript-sequencing. NG, mNeonGreen. NLS, nuclear localization signal. NES, nuclear export sequence.

### HDR-based EPHX2 nuclear import improves the cardiac function of *Lmna*-deficienct hearts

The above studies indicated that *Lmna* deficiency reduced cytoplasmic EPHX2 and increased nuclear EPHX2, which were both beneficiary to the heart (**Figure 6A**). To simultaneously promote both effects, a homology-directed repair (HDR) mechanism was established to knock-in NLS into endogenous *Ephx2* (**Figure 6B**), which harnessed the unique high efficiency of HDR in cardiomyocytes using AAV-delivered single-strand DNA as the donor in the Rosa^CAG-LSL-SpCas9-GFP^ mice[64]. Amplicon-sequencing analysis confirmed that the coding sequence of NLS was successfully inserted next to the start codon of *Ephx2* in a fraction of cells in the heart while the rest of cells remain either intact or undergoing non-homologous end jointing (NHEJ) (**Figure 6C**). Three sgRNAs were found to trigger this HDR reaction and the sgRNA with the highest total editing rate was selected for further analysis (**Figure S4A-B**).

**Figure 6.**
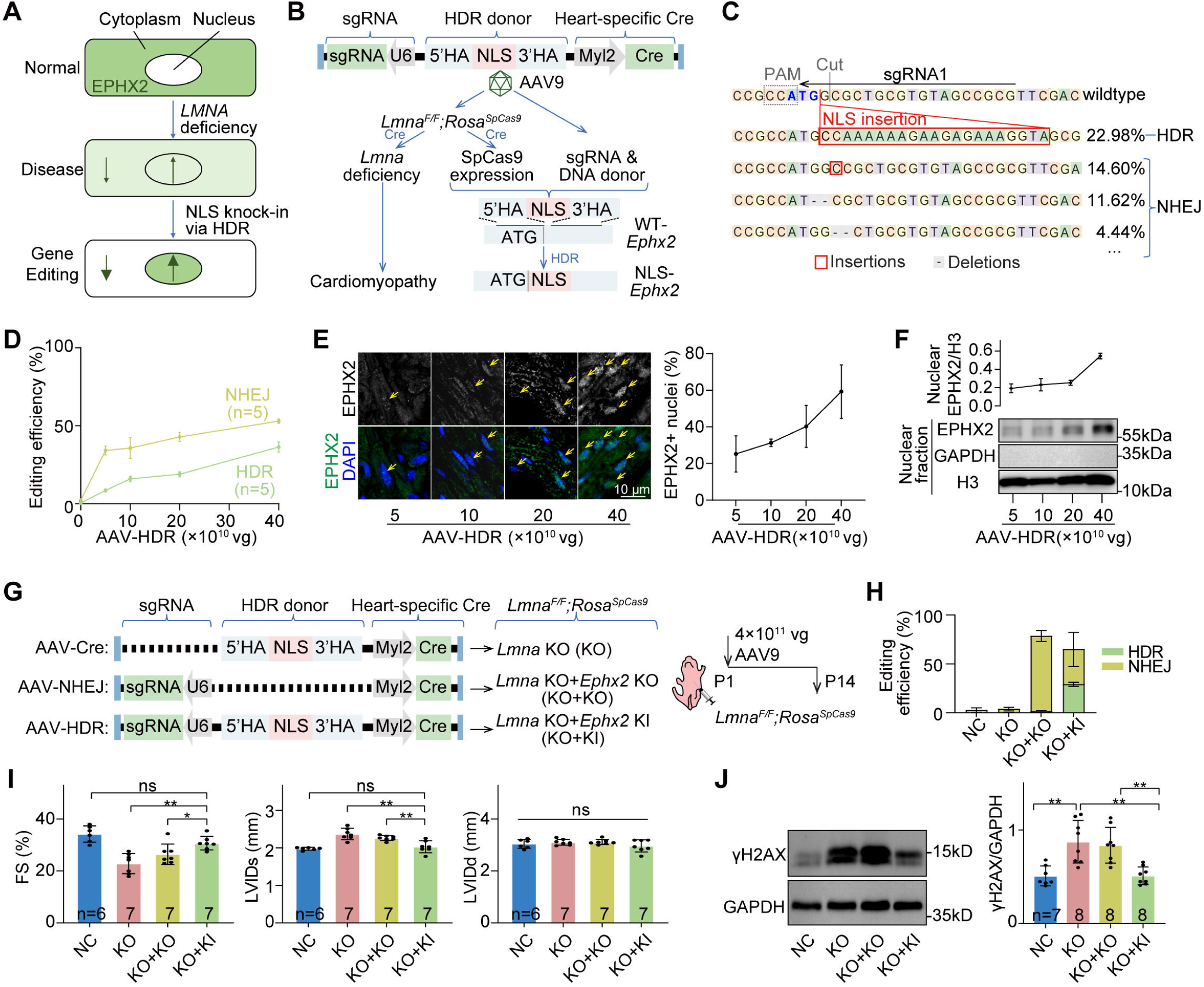
AAV-HDR system mediates nuclear localization of endogenous EPHX2 and alleviates *Lmna*-associated cardiac dysfunction. **A**, Diagram of gene therapy of NLS knock-in via HDR. **B**, Work flow of the AAV-HDR system. **C**, Example of targeted cDNA amplicon-sequencing results. The edited genomic regions were analyzed using CRISPResso2 software, and the editing efficiencies of HDR and NHEJ were calculated separately. **D**, AAV dose-dependent editing efficiency in mouse hearts evaluated by amplicon sequencing. **E**, Immunofluorescence images and quantification of EPHX2-positive nucleus in myocardium. n=3. **F**, Western blot images and quantification of nuclear EPHX2. n=3. **G**, Diagram of AAV-HDR and control AAVs, and outcomes in *Lmna^F/F^*; Rosa^CAG-LSL-Cas9-tdGFP^ mice after treatment. **H**, Editing efficiency of HDR and NHEJ-mediated gene edting after 4×10^11^ AAV treatment by amplicon sequencing. n=4. **I**, Echocardiogram analysis of AAV-HDR treated hearts. **J**, Western blot images and quantification of γH2AX in myocardium. *P* values were generated by Mann-Whitney *U* test. \**P*<0.05, \*\**P* < 0.01, \*\*\**P* < 0.001, mean ± SD. n represents numbers of mice. HDR, homology-directed repair. NHEJ, non-homologous end-joining. HA, homology arm. Cre, Cyclization recombination. LSL, loxP-stop-loxP. NLS, nuclear localization signal. KO, knock-out. KI, knock-in. FS, fraction shortening. LVIDd, left ventricular internal dimension in diastole. LVIDs, left ventricular internal dimension in systole.

Serial doses of AAVs were next applied to the heart and dose-dependent gene editing of HDR and NHEJ was detected with upper limits of ∼38% and ∼55% editing rates (**Figure 6D**). Since NHEJ leads to *Ephx2* knockout, which is beneficiary to the heart, the concurrent NHEJ unlikely counteracted the effect of HDR. By immunofluorescence and nuclear fractionation analyses, the nuclear translocation of endogenous EPHX2 by HDR was also observed in a dose-dependent manner (**Figure 6E-F**). Strikingly, HDR-based EPHX2 relocation was detected in up to 60% cardiomyocytes (**Figure S4C**). This higher protein translocation rate than HDR editing rate is presumably because most cardiomyocytes are tetraploid and HDR in only 1/4 *Ephx2* alleles is sufficient to trigger detectable translocation at the protein level.

Next, the AAV-HDR vector was compared to control vectors that lack either sgRNA (AAV-Cre, *Lmna* KO) or donor DNA (AAV-NHEJ, *Lmna* KO + *Ephx2* KO) to evaluate their effects on the heart with *Lmna* deficiency (**Figure 6G**). High and comparable editing rates were confirmed by amplicon sequencing in both AAV-HDR and AAV-NHEJ groups (**Figure 6H**). Echocardiographic analysis revealed pronounced mitigation of cardiac dysfunction by AAV-HDR compared with AAV-NHEJ (**Figure 6I**). The level of γH2AX was markedly attenuated in the HDR group, agreeing to the favorable impact of nuclear EPHX2 on DNA damage (**Figure 6J**).

## Discussion

Systemic identification and manipulation of key subcellular translocations in cardiac pathogenesis remain challenging in vivo. To solve this problem, here proximity proteomics was combined with HDR gene editing to build a platform to both discover and engineer endogenous protein localization specifically in cardiomyocytes in mice. Using *LMNA*-related cardiomyopathy as an example, this study found EPHX2 nuclear translocation as a novel target that could potentially be harnessed to mitigate *LMNA*-related cardiac dysfunction.

*LMNA*-related cardiomyopathy is a difficult indication for gene therapy due to the high heterogeneity of disease-causing variants and the complicated pathological mechanisms. *LMNA* gene supplementation[24, 46], *SUN1* gene suppression[24, 25] and base editing-based variant correction[27, 65] strategies are each suitable for only a small fraction of *LMNA* variants. Without the potential to expand the spectrum of applicable indications, the costs of these therapeutics are also difficult to afford. Therefore, identifying broad-spectrum therapeutic targets like EPHX2, which holds the promise to expand gene therapy from rare diseases to common diseases, are critical to accelerate translational medicine at the next stage.

EPHX2 was classically a cytoplasmic hydrolase that aggravates many cardiovascular diseases. Its hydrolase-independent function was implicated previously[45] but poorly studied in the heart. Localization changes of EPHX2 in cardiomyocytes upon *Lmna* deficiency generated a unique opportunity to uncover its ectopic function in nuclei. This is also an important example that subcellular translocation changes naturally involve two sides of the mechanism, namely the loss of functions in the original compartment and the generation of ectopic functions in the new compartment. Systemic investigations of both sides are necessary to fully understand the meaning of a given change of subcellular localization.

HDR gene editing usually exhibits low efficiency in postmitotic cells. However, cardiomyocytes, particularly at the perinatal stage, demonstrate high HDR rates[64, 66] presumably because of their robust DNA replication and DNA repair activity. DNA replication without cytokinesis leads to tetraploidy in cardiomyocytes that amplifies the outcome of HDR-based protein translocation. For example, HDR-based NLS knockin in *Ephx2* resulted in ∼30% HDR rate but detectable EPHX2 nuclear import in ∼60% cardiomyocytes, which is likely because gene editing in 1/4 alleles is sufficient to trigger detectable nuclear localization in cardiomyocytes.

It is critical to notice that somatic HDR is always mixed with NHEJ in the cell population. Although methods to enhance the HDR-NHEJ ratio are emerging[67, 68], complete elimination of NHEJ is unlikely to achieve in vivo. Therefore, in vivo HDR should be restricted to applications where the outcomes of HDR and NHEJ are not contradictory. For example, in this study, NHEJ-based *Ephx2* silencing is beneficiary to the heart. The HDR editing of NLS knock-in added additional benefits while keeping the NHEJ effect unaltered.

## Acknowledgements

The authors acknowledge Dr Yixian Zheng for providing the *Lmna^F/F^* mice and PackGene Biotech for AAV production.

## Nonstandard Abbreviations and Acronyms

AAV: adeno-associated virus
CRISPR: clustered regularly interspaced short palindromic repeats
Cas9: CRISPR-associated protein 9
EET: epoxyeicosatrienoic acid
EPHX2: epoxide hydrolase 2
HDR: homology-directed repair
LMNA: lamin-A/C
NES: nuclear export signal
NLS: nuclear localization signal

## References

1. Koch, A.J. and J.M. Holaska, Emerin in health and disease. Semin Cell Dev Biol, 2014. 29: p. 95–106.

2. Le Dour, C., et al., Actin-microtubule cytoskeletal interplay mediated by MRTF-A/SRF signaling promotes dilated cardiomyopathy caused by LMNA mutations. Nat Commun, 2022. 13(1): p. 7886.

3. Liu, B., et al., Resveratrol rescues SIRT1-dependent adult stem cell decline and alleviates progeroid features in laminopathy-based progeria. Cell Metab, 2012. 16(6): p. 738–50.

4. Conforti, F., et al., Molecular Pathways: Anticancer Activity by Inhibition of Nucleocytoplasmic Shuttling. Clin Cancer Res, 2015. 21(20): p. 4508–13.

5. Aebi, U., et al., The nuclear lamina is a meshwork of intermediate-type filaments. Nature, 1986. 323(6088): p. 560–4.

6. Turgay, Y., et al., The molecular architecture of lamins in somatic cells. Nature, 2017. 543(7644): p. 261–264.

7. Broers, J.L., et al., Decreased mechanical stiffness in LMNA-/- cells is caused by defective nucleo-cytoskeletal integrity: implications for the development of laminopathies. Hum Mol Genet, 2004. 13(21): p. 2567–80.

8. Dahl, K.N., et al., The nuclear envelope lamina network has elasticity and a compressibility limit suggestive of a molecular shock absorber. J Cell Sci, 2004. 117(Pt 20): p. 4779–86.

9. Swift, J., et al., Nuclear lamin-A scales with tissue stiffness and enhances matrix-directed differentiation. Science, 2013. 341(6149): p. 1240104.

10. Earle, A.J., et al., Mutant lamins cause nuclear envelope rupture and DNA damage in skeletal muscle cells. Nat Mater, 2020. 19(4): p. 464–473.

11. Cho, S., et al., Mechanosensing by the Lamina Protects against Nuclear Rupture, DNA Damage, and Cell-Cycle Arrest. Dev Cell, 2019. 49(6): p. 920–935.e5.

12. En, A., et al., Pervasive nuclear envelope ruptures precede ECM signaling and disease onset without activating cGAS-STING in Lamin-cardiomyopathy mice. Cell Rep, 2024. 43(6): p. 114284.

13. Ho, C.Y., et al., Lamin A/C and emerin regulate MKL1-SRF activity by modulating actin dynamics. Nature, 2013. 497(7450): p. 507–11.

14. Guo, Y., et al., Concentration-dependent lamin assembly and its roles in the localization of other nuclear proteins. Mol Biol Cell, 2014. 25(8): p. 1287–97.

15. Ghosh, S., B. Liu, and Z. Zhou, Resveratrol activates SIRT1 in a Lamin A-dependent manner. Cell Cycle, 2013. 12(6): p. 872–6.

16. Qiu, H., et al., Lamin A/C deficiency-mediated ROS elevation contributes to pathogenic phenotypes of dilated cardiomyopathy in iPSC model. Nat Commun, 2024. 15(1): p. 7000.

17. Cheedipudi, S.M., S. Asghar, and A.J. Marian, Genetic Ablation of the DNA Damage Response Pathway Attenuates Lamin-Associated Dilated Cardiomyopathy in Mice. JACC Basic Transl Sci, 2022. 7(12): p. 1232–1245.

18. Roux, K.J., et al., A promiscuous biotin ligase fusion protein identifies proximal and interacting proteins in mammalian cells. J Cell Biol, 2012. 196(6): p. 801–10.

19. May, D.G., et al., Comparative Application of BioID and TurboID for Protein-Proximity Biotinylation. Cells, 2020. 9(5).

20. Hershberger, R.E., D.J. Hedges, and A. Morales, Dilated cardiomyopathy: the complexity of a diverse genetic architecture. Nat Rev Cardiol, 2013. 10(9): p. 531–47.

21. Hasselberg, N.E., et al., Lamin A/C cardiomyopathy: young onset, high penetrance, and frequent need for heart transplantation. Eur Heart J, 2018. 39(10): p. 853–860.

22. Paldino, A., et al., Prognostic Prediction of Genotype vs Phenotype in Genetic Cardiomyopathies. J Am Coll Cardiol, 2022. 80(21): p. 1981–1994.

23. Garcia-Pavia, P., et al., REALM-DCM: A Phase 3, Multinational, Randomized, Placebo-Controlled Trial of ARRY-371797 in Patients With Symptomatic LMNA-Related Dilated Cardiomyopathy. Circ Heart Fail, 2024. 17(7): p. e011548.

24. Tan, C.Y., et al., Systematic in vivo candidate evaluation uncovers therapeutic targets for LMNA dilated cardiomyopathy and risk of Lamin A toxicity. J Transl Med, 2023. 21(1): p. 690.

25. Chai, R.J., et al., Disrupting the LINC complex by AAV mediated gene transduction prevents progression of Lamin induced cardiomyopathy. Nat Commun, 2021. 12(1): p. 4722.

26. Chen, C.Y., et al., Accumulation of the inner nuclear envelope protein Sun1 is pathogenic in progeric and dystrophic laminopathies. Cell, 2012. 149(3): p. 565–77.

27. Caravia, X.M., et al., Precise gene editing of pathogenic Lamin A mutations corrects cardiac disease. Proc Natl Acad Sci U S A, 2025. 122(43): p. e2515267122.

28. Wang, Z., et al., LMNA-related cardiomyopathy: From molecular pathology to cardiac gene therapy. J Adv Res, 2025. 77: p. 443–464.

29. Yang, L., et al., The LMNA p.R541C mutation causes dilated cardiomyopathy in human and mice. Int J Cardiol, 2022. 363: p. 149–158.

30. Kato, K., et al., LMNA Missense Mutation Causes Nonsense-Mediated mRNA Decay and Severe Dilated Cardiomyopathy. Circ Genom Precis Med, 2020. 13(5): p. 435–443.

31. Gupta, P., et al., Genetic and ultrastructural studies in dilated cardiomyopathy patients: a large deletion in the lamin A/C gene is associated with cardiomyocyte nuclear envelope disruption. Basic Res Cardiol, 2010. 105(3): p. 365–77.

32. Bhaskaran, A., et al., Location of LMNA Variants and Clinical Outcomes in Cardiomyopathy. JAMA Cardiol, 2025. 10(9): p. 896–903.

33. Spector, A.A., et al., Epoxyeicosatrienoic acids (EETs): metabolism and biochemical function. Prog Lipid Res, 2004. 43(1): p. 55–90.

34. Imig, J.D., L. Cervenka, and J. Neckar, Epoxylipids and soluble epoxide hydrolase in heart diseases. Biochem Pharmacol, 2022. 195: p. 114866.

35. Imig, J.D., Epoxides and soluble epoxide hydrolase in cardiovascular physiology. Physiol Rev, 2012. 92(1): p. 101–30.

36. Xu, X., X.A. Zhang, and D.W. Wang, The roles of CYP450 epoxygenases and metabolites, epoxyeicosatrienoic acids, in cardiovascular and malignant diseases. Adv Drug Deliv Rev, 2011. 63(8): p. 597–609.

37. Qiu, H., et al., Soluble epoxide hydrolase inhibitors and heart failure. Cardiovasc Ther, 2011. 29(2): p. 99–111.

38. Ma, K., et al., Therapeutic and Prognostic Significance of Arachidonic Acid in Heart Failure. Circ Res, 2022. 130(7): p. 1056–1071.

39. Gui, Y., et al., Soluble epoxide hydrolase inhibitors improve angiogenic function of endothelial progenitor cells via ERK/p38-mediated miR-126 upregulation in myocardial infarction mice after exercise. Exp Cell Res, 2020. 397(2): p. 112360.

40. Wang, Y.X., et al., Soluble epoxide hydrolase in atherosclerosis. Curr Atheroscler Rep, 2010. 12(3): p. 174–83.

41. Wang, Q., et al., Soluble epoxide hydrolase is involved in the development of atherosclerosis and arterial neointima formation by regulating smooth muscle cell migration. Am J Physiol Heart Circ Physiol, 2015. 309(11): p. H1894–903.

42. He, W., et al., Deletion of soluble epoxide hydrolase suppressed chronic kidney disease-related vascular calcification by restoring Sirtuin 3 expression. Cell Death Dis, 2021. 12(11): p. 992.

43. Cai, Y., et al., Diabetic vascular calcification inhibited by soluble epoxide hydrolase gene deletion via regressing NID2-mediated IGF2-ERK1/2 signaling pathway. Chin Med J (Engl), 2025. 138(20): p. 2657–2668.

44. Rawal, S., et al., Differential subcellular distribution and colocalization of the microsomal and soluble epoxide hydrolases in cultured neonatal rat brain cortical astrocytes. J Neurosci Res, 2009. 87(1): p. 218–27.

45. Lien, C.C., et al., The phosphatase activity of soluble epoxide hydrolase regulates ATP-binding cassette transporter-A1-dependent cholesterol efflux. J Cell Mol Med, 2019. 23(10): p. 6611–6621.

46. Sun, Y., et al., Non-Cell-Autonomous Cardiomyocyte Regulation Complicates Gene Supplementation Therapy for Lmna-Associated Cardiac Defects in Mice. JACC Basic Transl Sci, 2024. 9(11): p. 1308–1325.

47. Agah, R., et al., Gene recombination in postmitotic cells. Targeted expression of Cre recombinase provokes cardiac-restricted, site-specific rearrangement in adult ventricular muscle in vivo. J Clin Invest, 1997. 100(1): p. 169–79.

48. Kim, Y. and Y. Zheng, Generation and characterization of a conditional deletion allele for Lmna in mice. Biochem Biophys Res Commun, 2013. 440(1): p. 8–13.

49. Kim, D.I., et al., An improved smaller biotin ligase for BioID proximity labeling. Mol Biol Cell, 2016. 27(8): p. 1188–96.

50. Guo, Y., et al., Sarcomeres regulate murine cardiomyocyte maturation through MRTF-SRF signaling. Proc Natl Acad Sci U S A, 2021. 118(2).

51. Yang, L., et al., MicroRNA-122-Mediated Liver Detargeting Enhances the Tissue Specificity of Cardiac Genome Editing. Circulation, 2024. 149(22): p. 1778–1781.

52. Guo, Y., et al., Analysis of Cardiac Myocyte Maturation Using CASAAV, a Platform for Rapid Dissection of Cardiac Myocyte Gene Function In Vivo. Circ Res, 2017. 120(12): p. 1874–1888.

53. Guo, C.T., et al., In vivo proximity proteomics uncovers palmdelphin (PALMD) as a Z-disc-associated mitigator of isoproterenol-induced cardiac injury. Acta Pharmacol Sin, 2024. 45(12): p. 2540–2552.

54. von Gise, A., et al., YAP1, the nuclear target of Hippo signaling, stimulates heart growth through cardiomyocyte proliferation but not hypertrophy. Proc Natl Acad Sci U S A, 2012. 109(7): p. 2394–9.

55. Sullivan, T., et al., Loss of A-type lamin expression compromises nuclear envelope integrity leading to muscular dystrophy. J Cell Biol, 1999. 147(5): p. 913–20.

56. Ødum, M.T., et al., DeepLoc 2.1: multi-label membrane protein type prediction using protein language models. Nucleic Acids Res, 2024. 52(W1): p. W215–w220.

57. Horton, P., et al., WoLF PSORT: protein localization predictor. Nucleic Acids Res, 2007. 35(Web Server issue): p. W585–7.

58. Chou, K.C. and H.B. Shen, Cell-PLoc: a package of Web servers for predicting subcellular localization of proteins in various organisms. Nat Protoc, 2008. 3(2): p. 153–62.

59. Kosugi, S., et al., Systematic identification of cell cycle-dependent yeast nucleocytoplasmic shuttling proteins by prediction of composite motifs. Proc Natl Acad Sci U S A, 2009. 106(25): p. 10171–6.

60. Brameier, M., A. Krings, and R.M. MacCallum, NucPred--predicting nuclear localization of proteins. Bioinformatics, 2007. 23(9): p. 1159–60.

61. Auguste, G., et al., BET bromodomain inhibition attenuates cardiac phenotype in myocyte-specific lamin A/C-deficient mice. J Clin Invest, 2020. 130(9): p. 4740–4758.

62. VanDusen, N.J., et al., CASAAV: A CRISPR-Based Platform for Rapid Dissection of Gene Function In Vivo. Curr Protoc Mol Biol, 2017. 120: p. 31.11.1–31.11.14.

63. Pinot, F., et al., Molecular and biochemical evidence for the involvement of the Asp-333-His-523 pair in the catalytic mechanism of soluble epoxide hydrolase. J Biol Chem, 1995. 270(14): p. 7968–74.

64. Zheng, Y., et al., Efficient In Vivo Homology-Directed Repair Within Cardiomyocytes. Circulation, 2022. 145(10): p. 787–789.

65. Yang, L., et al., Adenine base editor-based correction of the cardiac pathogenic Lmna c.1621C > T mutation in murine hearts. J Cell Mol Med, 2024. 28(4): p. e18145.

66. Ishizu, T., et al., Targeted Genome Replacement via Homology-directed Repair in Non-dividing Cardiomyocytes. Sci Rep, 2017. 7(1): p. 9363.

67. Yeh, C.D., C.D. Richardson, and J.E. Corn, Advances in genome editing through control of DNA repair pathways. Nat Cell Biol, 2019. 21(12): p. 1468–1478.

68. Fu, Y.W., et al., Dynamics and competition of CRISPR-Cas9 ribonucleoproteins and AAV donor-mediated NHEJ, MMEJ and HDR editing. Nucleic Acids Res, 2021. 49(2): p. 969–985.

