## Supplementary information for "*Lmna* deficiency promotes EPHX2 nuclear translocation to ameliorate cardiac dysfunction in mice"

### 1. Supplementary Figure

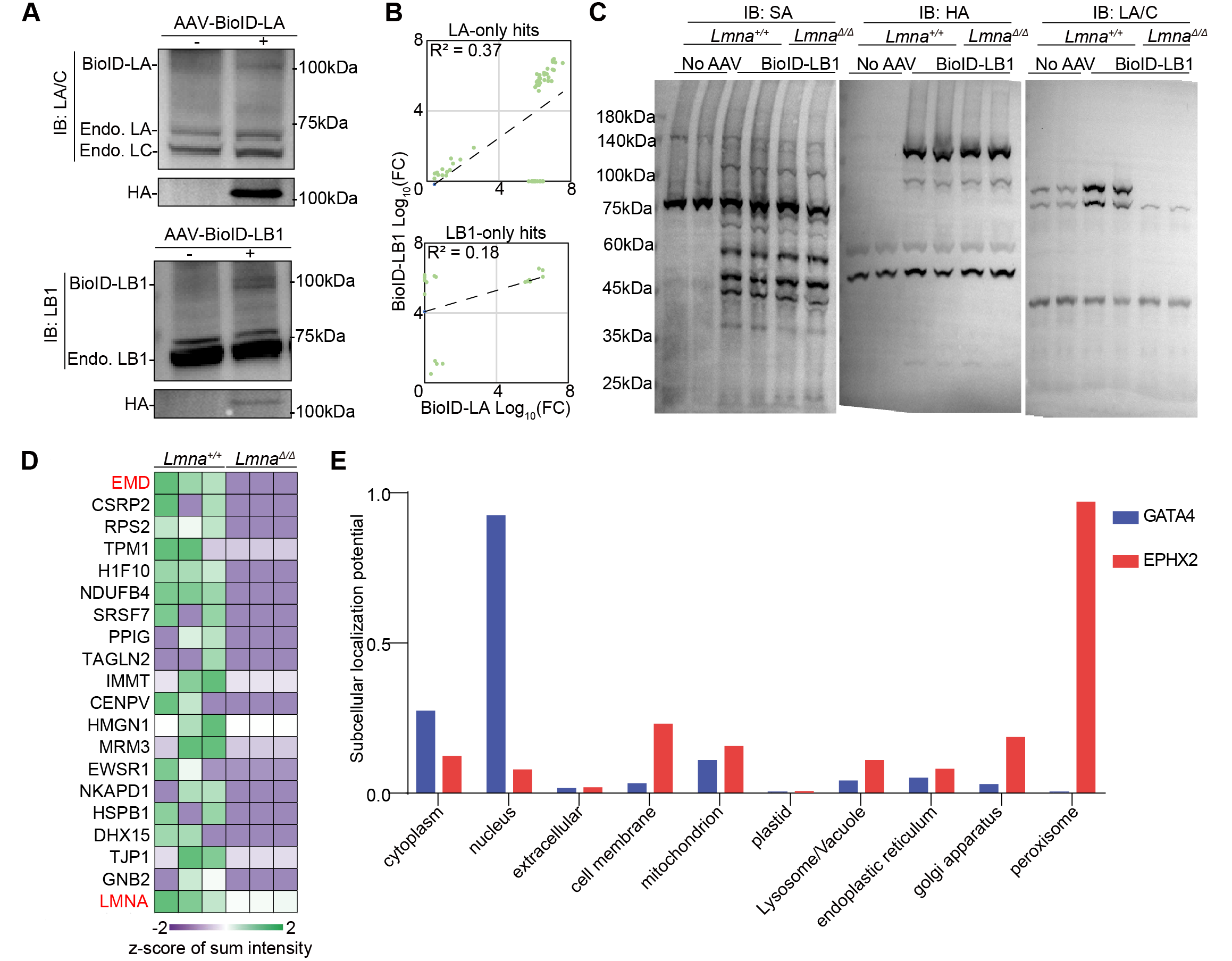

**Figure S1. In vivo proximity proteomics in *Lmna^Δ/Δ^* mice and subcellular localization prediction of EPHX2. A**, Western blotting of cardiac lysates. IB, immunoblottingEndo., endogenous. **B**, Correlation analysis of significantly enriched proteins. **C**, Western blotting of heart tissues from BioID-LB1-treated *Lmna^Δ/Δ^* mice. **D**, Heatmap of significantly downregulated proteins in *Lmna^Δ/Δ^* mouse hearts. **E**, Probability of predicted subcelullar localizations and membrane association by DeepLoc 2.1.

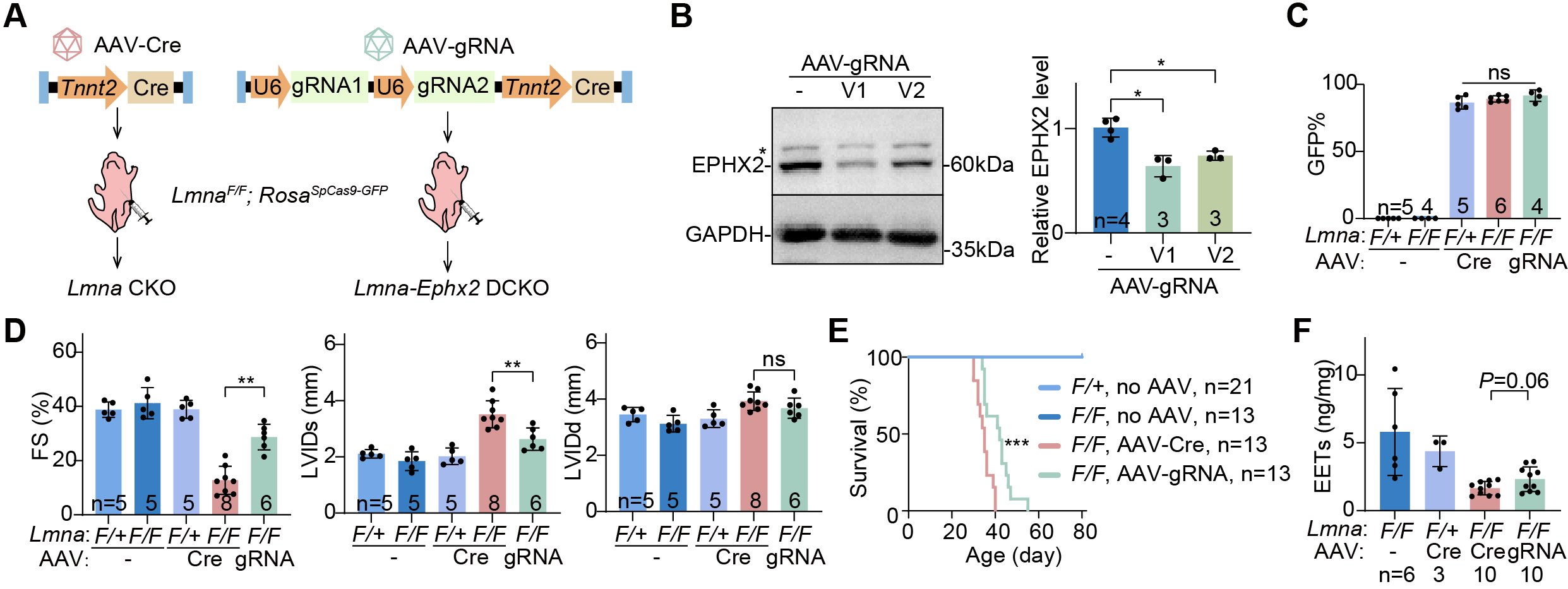

**Figure S2. CASAAV-based cardiomyocyte-specific EPHX2 depletion alleviates *Lmna*-deficient dysfunction. A**, a diagram showing the workflow of CASAAV. **B**, Western blot analysis of CASAAV-treated cardiac tissues. **C**, Quantification of GFP-positive cardiomyocytes in CASAAV-treated hearts. **D**, Echocardiogram analysis of CASAAV-treated hearts. **E**, Survival curve with the log-rank test between AAV-Cre and AAV-gRNA treated *Lmna^F/F^*; *Rosa^SpCas9-GFP^* mice, ****P*<0.05. **F,** EET quantification by ELISA. Mann-Whitney *U* test, **P* < 0.05, ***P* < 0.01, mean±SD. In **B-F**, n indicates animal numbers.

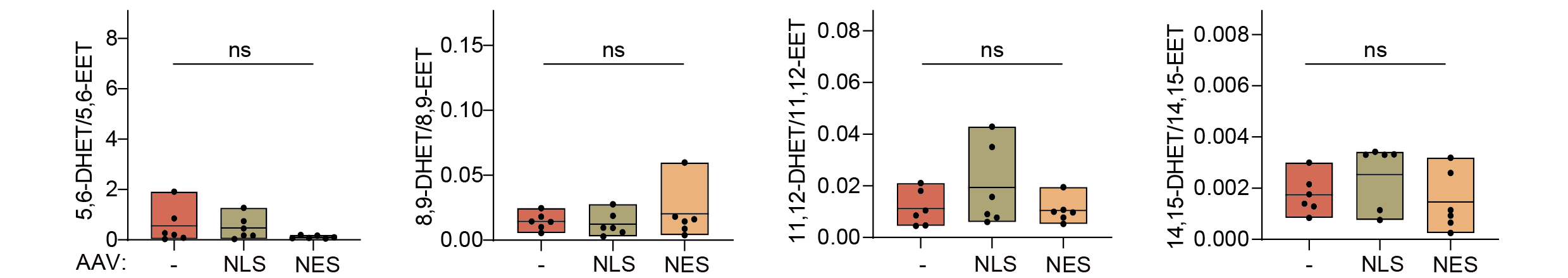

**Figure S3. DHET/EET levels in CKO mice after AAV treatment.** DHET/EET in hearts by ultra performance liquid chromatography-tandem mass spectrometry (LC–MS/MS)-based lipidomic. 6 animals per group. The Kruskal-Wallis *H* test was used for statistical analysis. NLS, nuclear localization sequence. NES, nuclear export sequence. ns, not significant.

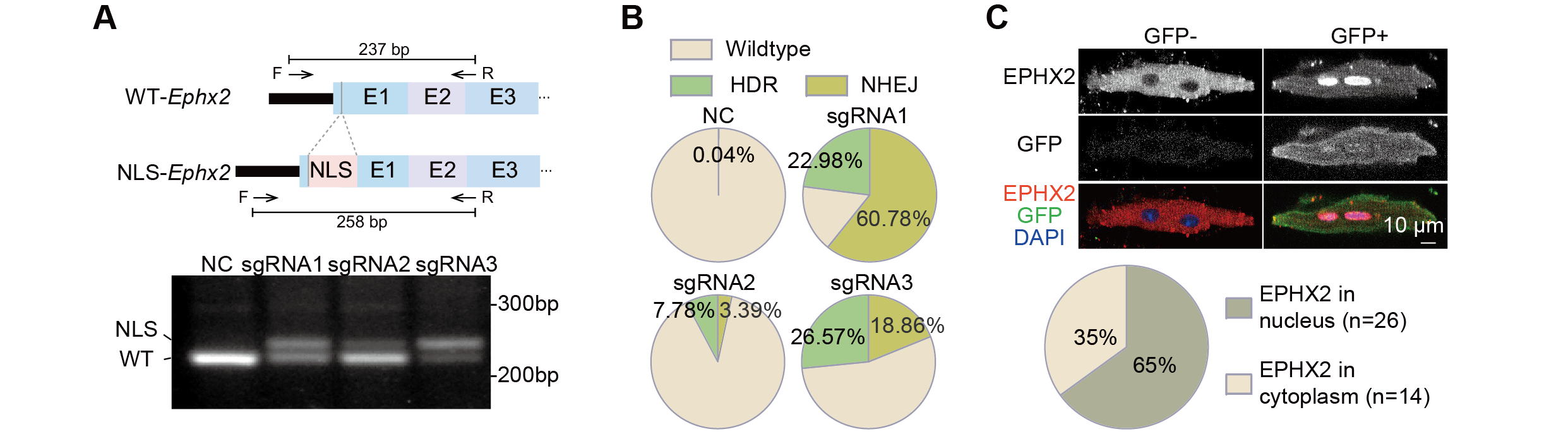

**Figure S4. Validation of the AAV-HDR system. A,** Schematic diagram of *Ephx2* gene RT-PCR and DNA agarose gel electrophoresis image. **B,** Amplicon-sequencing analysis of gene editing ratio by HDR and NHEJ of all 3 sgRNAs. **C,** immunofluorescence analysis and quantification of EPHX2 nuclear translocation following isolation of single cardiomyocytes.

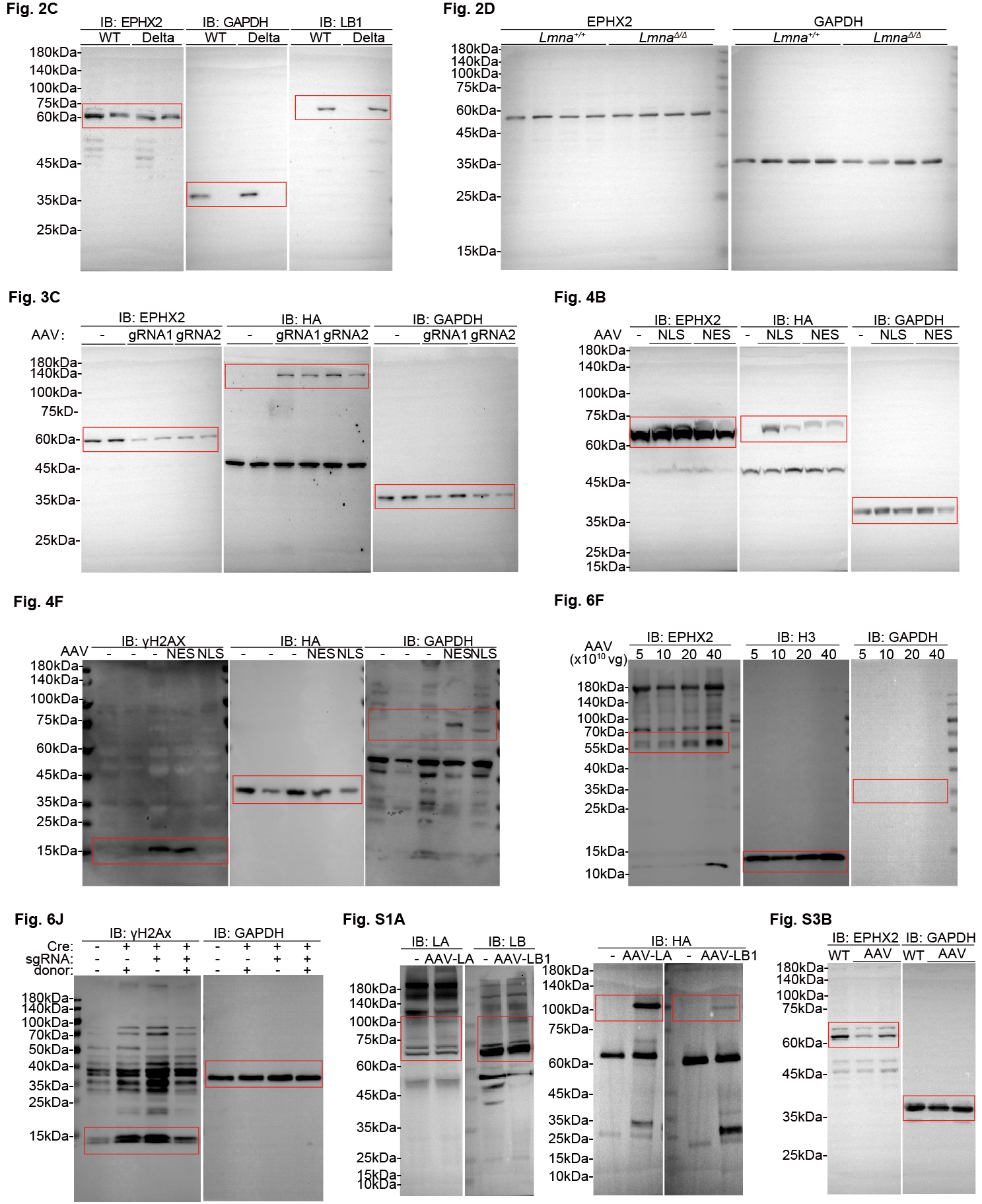

**Figure S5**. **Full- length raw western blots for each panel.**

### 2. Supplementary Table

#### Table S1. Genome editing

| **Gene** | **sgRNA** |
| --- | --- |
| sgRNA targeting *Ephx2* by SaCas9 | sgRNA1: 5’-GCCAACACTTTGACTTCCTGA-3’ sgRNA2: 5’-GCAATCCAAATGACGTCAGCC-3’ |
| sgRNA targeting *Ephx2* by CASAAV | Version1: 5’-TGGTAACCATCCCCATGTCA-3’  5’-AGTCATCTAGGAAAACAACC-3’ Version2: 5’-ACCAACAACTGGCTGGACGA-3’  5’-AGTAAAAAATTGTAGATCTG-3’ |
| sgRNA targeting *Ephx2* for HDR | sgRNA1: 5’-CGCGGCTACACGCAGCGCCA-3’  sgRNA2: 5’-GGCTACACGCAGCGCCATGG-3’ sgRNA3: 5’-CACTCCGTCAAGGTCGAACG-3’ |
| DNA donor for NLS knock-in* | 5’-GACCCTTTCTGTTCTCAGGGCAACACCCCCACATTTTCAGCCACTTTCCATTTCTTAGGTGTGAATCTCAGAAAACACAAACTGCCCCAGACAAGACTTGAGGCAAGAAGAGGAAGTGTCTGAAAGACTCACAAAAATTAACCCACAGTAGGGCCATTTCCCACCACCTAACACGCAGCAACCAGAGCAAAGGAAGAAGCCAATTCCCGGTTTCAGTAAGGGAGGCAGTCTTTCACAACAGCCTTGGACTAAACGGTGGATTTGTTAATTAACTCATTAGTAGAGGAGGTTTCTCTTTTTTCTCTTAACTAGGGAGAGCACTGGGCCAAGGGGCAGGAAGGTTCCAGAAACCTCCAAGAAGCCCAGCTCACTTTTAAACAATACAGTGTAGAAAAAGGAAACAGACTCGATTAATTTCTTTTAATCCTCAAGAGAAAAAGTATTCTGGGTTAGCACACCAAGACAGGAGGCTGTCTTGCATTCCAGAAACACTGCTAGAAGAAGCAGGCAGATGTGAGGGGGTTGCAAAGATGTGGACCACATTTAGAGAAGAGGGTCCACAGATCCCAACATACATGGGATATGCCAATCAAGACCAAATACTGAAGGAGAAAATGTTTAGAATATTCCGTCTCCTCCAACCTGCCTCCTCAAGACCCCTCTCTCCCCATACCAAGCCTGTTTCCAAACGGTCAAAGTCTGAGAATTGGGGAGCCTTGGCAGGGTTTCTAGTCCTTAGGCTAAGGAGGAAAAAGTGGCCAAGACCAGGATTGACACTTGGGTGGGACATCCAGCAAGAAGCTCCGGGCTAGGCAGCTGATCAGTAGGCAGGGCAGTGTCAGTACTGGGCGGGGCGGGGCGGGGCGGGGCGGGGCGAGGGGCGGTGCTGAGATTGGCACGACCCTAATCTTAGGTTCCCACAGCCAGCCTCCCCACTCTAGGCCACAGCCTTCCAGCTTCGTGTCTGTGTCAGCTTGACGCTGCAGCCCGGGCCGCCATG**CCAAAAAAGAAGAGAAAGGTA**GCGCTGCGTGTAGCCGCGTTCGACCTTGACGGAGTGCTGGCCCTCCCCTCTATCGCCGGGGCTTTCCGCCGCAGCGAAGAGGCCCTGGCACTGCCTAGGTAAGGGGAGCCGCGCTGGGTCAGTGTCCCGCCTGGTGCCCTGTCCATACGTCCAGCCCAGGTTTCAAATTGCTTTCCAGGTGAGCCTGGGCAAGATCAAAATCATTACCTGGTGTTAAAATGCCAGCGTGCACGAGTAGATCTATTGCCCTTCTAAATGCTAAGTAATTAAAACTTGCTTTAAAAAAAAACTCCTTCATGAAAAGAGGCGTTATTACCATTTGCTGCCGGATTAATGCTACTGAACTGGGTCTTTAAAATACCCGCCGCGATTTACAAGAAAGTTCAGTCCTAGGCCAGCGTTATAGGAGAACTAGTGCAAGAGCAGGTTACAGAAATGTCCCACAGCCCAGGGAAACGGGAACCTGGGAGGCAGTCACTAATATCAGAAAGTAGTAGCTCTGATTCAGACCGAGGAGGGAGGAAGCAGGAAACAGTAGAAAATGTCTGGAATAAATATATCAAAATATCCAGACATGGGACTGGAGAGACGACTAAAGTTAAGAAAGTGCTCTTTCGAGCCACCATGTGGTTTCTGGGATTTGAACTCAGGACCTTTGGAAGAGCAGGGCAATAGTGGCACACGCCTTTAATCCCAGCACTTGGGAGGCAGAGGCAGGCGAATTTCTAAGTTTGAGGCCAGCTTGGTCTACAGAGTGAGTTCCAGGACAGCCAGGGCTACACAGAGAAACCCTGTCTCGAAAAAAAAAAAAAAAGCAAAACACAAAAACAAACAAGAAAGAAAGAAAGAGAAAGAAAGAAAGAAAGAAAGAAAGAAAGAAAGAAAGAAAGAAAGAAAGAAAGAAAGGTGTTCTTGCTGAACTTTTGAGGCCCAGCAACCAAGTCAGGCGGCTCCGGCGGATCTGACTCCCCCTTCTGACGTCTTATGATACCTGTATGCA-3’ |

*NLS sequence in bold.

#### Table S2. Genotyping

| **Genotype** | **Primer F** | **Primer R** |
| --- | --- | --- |
| *Lmna^△^* | 5’-AAGCTACAGGAAGTC  CTCCA-3’ | 5’-TGATGCTTCTTGCT  GCCATA-3’ |
| *Lmna^Flox^* | 5’-AACCCAGCCTCAGAAAC  TGGTGGATG-3’ | 5’-GACAGCTCTCCTCTGAA  GTGCTTGGA-3’ |
| *Cre* | 5’-GCCATAGGCTACGGTG  TAAAAG-3’ | 5’-ATAATCGCGAACA  TCTTCAGGT-3’ |

#### Table S3. RT-qPCR primers

| **Gene** | **Primer F** | **Primer R** |
| --- | --- | --- |
| *Ephx2* | 5’-AAGCTACAGGAAGTCCTCCA-3’ | 5’-TGATGCTTCTTGCTGCCATA-3’ |
| *Gapdh* | 5’-TGACCACAGTCCATGCCATC-3’ | 5’-GACGGACACATTGGGGGTAG-3’ |

#### Table S4. Amplicon sequencing primers*

| **Application** | **Barcoded forward primers** | **Barcoded reverse primers** |
| --- | --- | --- |
| Amplicon sequencing for *Ephx2* gene site1 | 5’-AATGATACGGCGACCACCGAGATCTACACAGGCGAAGACACTCTTTCCCTACACGACGCTCTTCCGATCTGGCAGAGCTCTCTCTATAGC-3’ | 5’-CAAGCAGAAGACGGCATACGAGATTTCTGAATGTGACTGGAGTTCAGACGTGTGCTCTTCCGATCTTTTCCCCGTGAAAGGGAGTA-3’ |
|  | 5’-AATGATACGGCGACCACCGAGATCTACACTAATCTTAACACTCTTTCCCTACACGACGCTCTTCCGATCTGGCAGAGCTCTCTCTATAGC-3’ | 5’-CAAGCAGAAGACGGCATACGAGATACGAATTCGTGACTGGAGTTCAGACGTGTGCTCTTCCGATCTTTTCCCCGTGAAAGGGAGTA-3’ |
| Amplicon sequencing for *Ephx2* gene site2 | 5’-AATGATACGGCGACCACCGAGATCTACACCAGGACGTACACTCTTTCCCTACACGACGCTCTTCCGATCTAGCCATGTCCAATCTGGATG-3’ | 5’-CAAGCAGAAGACGGCATACGAGATAGCTTCAGGTGACTGGAGTTCAGACGTGTGCTCTTCCGATCTTGACAATTCAGTGGTCCCTG-3’ |
|  | 5’-AATGATACGGCGACCACCGAGATCTACACGTACTGACACACTCTTTCCCTACACGACGCTCTTCCGATCTAGCCATGTCCAATCTGGATG-3’ | 5’-CAAGCAGAAGACGGCATACGAGATGCGCATTAGTGACTGGAGTTCAGACGTGTGCTCTTCCGATCTTGACAATTCAGTGGTCCCTG-3’ |
| Amplicon sequencing for *Ephx2* cDNA | 5’-AATGATACGGCGACCACCGAGATCTACACTATAGCCTACACTCTTTCCCTACACGACGCTCTTCCGATCTCTTCGTGTCTGTGTCAGCTT-3’ | 5’-CAAGCAGAAGACGGCATACGAGATCGAGTAATGTGACTGGAGTTCAGACGTGTGCTCTTCCGATCTGATTTTGATCTTGCCCAGGC-3’ |
|  |  | 5’-CAAGCAGAAGACGGCATACGAGATTCTCCGGAGTGACTGGAGTTCAGACGTGTGCTCTTCCGATCTGATTTTGATCTTGCCCAGGC-3’ |
|  |  | 5’-CAAGCAGAAGACGGCATACGAGATAATGAGCGGTGACTGGAGTTCAGACGTGTGCTCTTCCGATCTGATTTTGATCTTGCCCAGGC-3’ |
|  |  | 5’-CAAGCAGAAGACGGCATACGAGATGGAATCTCGTGACTGGAGTTCAGACGTGTGCTCTTCCGATCTGATTTTGATCTTGCCCAGGC-3’ |

*Barcodes in red and genome-matching sequences in green.

#### Table S5. Antibodies and dyes used in this study.

| **Antibody** | **Source** | **Vendor** | **Cat. No.** | **Application** |
| --- | --- | --- | --- | --- |
| EPHX2 Polyclonal antibody | Rabbit | Proteintech | 10833-1-AP | WB(1:2000); IF(1:1000) |
| HA-Tag (C29F4) Rabbit Monoclonal Antibody | Rabbit | Cell Signaling Technology | 3724 | WB (1:2000);  IF (1:1000) |
| Phospho-Histone H2A.X (Ser139) (20E3) Rabbit Monoclonal Antibody | Rabbit | Cell Signaling Technology | 9718 | WB (1:2000);  IF (1:1001) |
| Anti-GAPDH Mouse Monoclonal Antibody | Mouse | TransGen | HC301 | WB (1:5000) |
| Goat Anti-Mouse IgG (H+L), HRP Conjugate | Goat | TransGen | HS201 | WB (1:5000) |
| Goat Anti-Rabbit IgG (H+L), HRP Conjugate | Goat | TransGen | HS101 | WB (1:5000) |
| Donkey anti-Mouse IgG (H+L) Highly Cross-Adsorbed Secondary Antibody, Alexa Fluor™ 488 | Donkey | Thermo Scientific | A21202 | IF (1:1000) |
| Donkey anti-Rabbit IgG (H+L) Highly Cross-Adsorbed Secondary Antibody, Alexa Fluor™ 488 | Donkey | Thermo Scientific | A21206 | IF (1:1000) |
| Donkey anti-Mouse IgG (H+L) Highly Cross-Adsorbed Secondary Antibody, Alexa Fluor™ 555 | Donkey | Thermo Scientific | A31570 | IF (1:1000) |
| Donkey anti-Rabbit IgG (H+L) Highly Cross-Adsorbed Secondary Antibody, Alexa Fluor™ 555 | Donkey | Thermo Scientific | A31572 | IF (1:1000) |
| HRP-conjugated Streptavidin | NA | Sangon | D111054 | WB (1:500) |
| DAPI | NA | Thermo Scientific | 62248 | IF (1:1000) |

### 3. Supplementary Materials & Methods

#### RNA extraction and RT-qPCR

Total RNA was extracted from heart apex using the TransZol Up Plus RNA Kit (TransGen, Beijing, China, ER501-01). Genomic DNA removal and reverse transcription were conducted using TransScript II One-Step gDNA Removal and cDNA Synthesis SuperMix (TransGen, Beijing, China, AH311-03). Real-time PCR was performed using Perfect Start Green qPCR Super Mix (+DyeII) (TransGen, Beijing, China, AQ602-24) and analyzed by the AriaMx Real-Time PCR System (Agilent Technologies). See Table S3 for primer sequences.

#### Amplicon sequencing

cDNA reverse-transcribed from total RNA and genomic DNA were used for amplicon sequencing. Genomic DNA was extracted from heart tissues or cells with the TIANamp Genomic DNA Kit (DP304, TIANGEN). Target loci were amplified using Taq PCR MasterMix (KT211, TIANGEN) and purified with the TIANgel Purification Kit (DP219, TIANGEN). Amplicon sequencing was conducted on the Illumina NovaSeq 6000 platform at Novogene (Beijing, China). Sequencing results were analyzed via CRISPResso2. Primer sequences are provided in Table S4.

#### SMART-seq

SMART-seq was performed by Geekgene Technology (Beijing, China). Total RNA was extracted from isolated cardiomyocytes, and mNeonGreen-positive cardiomyocytes were sorted on a BD FACS Aria II SORP instrument with a 100 μm nozzle. KAPA HiFi HotStart Ready MIX (KAPA Biosystems, KK8504) and Ampure XP beads (Beckman, A63882) were used to construct and purify the library. Sequencing was operated on Illumina Novaseq 6000 platform using Paired-End PE150 method by Beijing Geek Gene Technology. FastQC (v0.12.1) software was used for quality control, and Trim Galore (v0.6.10) was used to remove low-quality reads and adapter sequences from the raw reads to obtain clean reads. The clean reads were then aligned to the mouse reference genome GRCm38 using STAR (v2.7.11b). Gene-level fragment counts were quantified using featureCounts in Subread (v2.1.1) based on GENCODE vM25 annotation. Differential expression analysis was performed using PyDESeq2 (v0.5.0). Genes with an absolute log2-fold change greater than 1 and an adjusted p-value (Benjamini-Hochberg) less than 0.05 were identified as significantly differentially expressed. Functional enrichment analysis was conducted using gseapy (v1.1.5) with MSigDB (v2024.1.Mm).

#### Cardiomyocyte isolation

Hearts of heparin-treated mice were extracted and cannulated onto a Langendorff perfusion apparatus. Perfusion buffer and collagenase II (Worthington, LS004177) were sequentially pumped into the heart to flush out blood and dissociate cardiomyocytes and equilibrate the heart at 37 °C. After 8 min of digestion, ventricles were cut from the digested heart, gently dissociated into single cardiomyocytes in 10% FBS/perfusion buffer and filtered through a 100 µm cell strainer to remove undigested tissues.

For immunostaining, cardiomyocytes were seeded on laminin-coated coverslips in Dulbecco’s modified Eagle medium (DMEM) supplemented with 10% fetal bovine serum (FBS) and cultured for 30 minutes. Cells were then fixed, permeabilized, and prepared for immunofluorescence analysis.

#### Immunofluorescence analysis

For immunofluorescence imaging of cardiomyocytes, cells were fixed with 4% paraformaldehyde, permeabilized and blocked in QuickBlock permeabilization and blocking solution (Beyotime, China, P0260). Then the cells were incubated with primary antibodies overnight at 4 °C and then incubated with secondary antibodies and/or dyes at room temperature for 1 h. Antibodies were diluted in QuickBlock immunofluorescence dilution solution (Beyotime, China, P0262/P0265) and washed 3 times by PBS. Then the cells were mounted with ProLong Diamond antifade mountant (Invitrogen, 36961).

For Immunofluorescence imaging of heart tissue, hearts were perfused with 4% paraformaldehyde for 10 min at room temperature. Subsequently, the hearts were placed in 10% sucrose solution at 4 °C overnight and then in 20% sucrose solution at 4 °C overnight for dehydration. Dehydrated samples were embedded in OCT (Sakura) and frozen at −20 °C. Heart sections (7 μm) were cut on a cryostat microtome (CM 1950, Leica, Germany). The following steps were the same as the cells. Confocal images were taken using a Leica TCS SP8 MP FLIM laser-scanning confocal microscope with a 40× objective or an Olympus FV3000 laser-scanning confocal microscope with a 40× objective. See Table S5 for antibody and dye information.

#### Generation and culture of MEFs

MEFs were isolated from E13.5 mouse embryos of fertile, age-appropriate mated mice. The day of vaginal plug observation was designated Embryonic day 0.5 (E0.5). Pregnant mice were sacrificed at E13.5 for embryo collection. Yolk sacs, heads and dark red internal organs were carefully removed from each embryo. Remaining embryonic tissues were digested with 0.25% trypsin (HUANKE, China, HK2109.11) to prepare MEF suspensions. After filtration through a 70 μm cell strainer, cells were resuspended and cultured in high-glucose DMEM (HUANKE, China, HK2109.07) supplemented with 10% FBS and 1×Penicillin-Streptomycin solution (Bio-Channel, China, BC-CE-007). P0 MEFs were passaged at a 1:5 ratio when reaching 70% confluency. An aseptic environment was maintained throughout the process.

#### Western blot

For total protein extraction, cells or heart tissues were homogenized in RIPA buffer (25 mM Tris pH7.4, 150 mM NaCl, 1% Triton X-100, 0.5% Na Deoxycholate, 0.1% SDS) supplemented with protease inhibitor cocktail and phosphatase inhibitor cocktail. For nuclear or cytoplasmic protein extraction and isolation, cells or hearts were lysed using Nuclear and Cytoplasmic Protein Extraction Kit (Beyotime, China, P0028). Lysates were denatured in 2× SDS sample buffer at 70 °C for 10 min, separated on a 10% Bis-Tris gradient gel (Sangon Biotech, C691101-0001), transferred to a PVDF membrane, and blocked by 4% BSA diluted in TBST. Primary antibodies were incubated with the membrane overnight at 4 °C. HRP-conjugated secondary antibodies were incubated for 1 h at room temperature. Antibodies diluted in TBST and washed 3 times by TBST after incubation. Chemiluminescence was detected using an Invitrogen iBright™ CL1500. See Table S5 for antibody information.

#### Echocardiography

Mice were anesthetized initially by 3% isoflurane before echocardiogram measurement and maintained asleep at 1%–1.5% isoflurane. Echocardiography was performed with a VINNO6n machine (VINNO Corporation, Suzhou, China) with a 23 MHz transducer. EF, FS, and LVID values were calculated by averaging results from five consecutive heart beats. Performers of echocardiography was blinded to mouse genotypes and treatments.
